# Effective cell membrane tension protects red blood cells against malaria invasion

**DOI:** 10.1101/2023.05.30.542792

**Authors:** Haleh Alimohamadi, Padmini Rangamani

## Abstract

A critical step in how malaria parasites invade red blood cells (RBCs) is the wrapping of the membrane around the egg-shaped merozoites. Recent experiments have revealed that RBCs can be protected from malaria invasion by high membrane tension. While cellular and biochemical aspects of parasite actomyosin motor forces during the malaria invasion have been well studied, the important role of the biophysical forces induced by the RBC membrane-cytoskeleton composite has not yet been fully understood. In this study, we use a theoretical model for lipid bilayer mechanics, cytoskeleton deformation, and membrane-merozoite interactions to systematically investigate the influence of effective RBC membrane tension, which includes contributions from the lipid bilayer tension, spontaneous tension, interfacial tension, and the resistance of cytoskeleton against shear deformation on the progression of membrane wrapping during the process of malaria invasion. Our model reveals that this effective membrane tension creates a wrapping energy barrier for a complete merozoite entry. We calculate the tension threshold required to impede the malaria invasion. We find that the tension threshold is a nonmonotonic function of spontaneous tension and undergoes a sharp transition from large to small values as the magnitude of interfacial tension increases. We also predict that the physical properties of the RBC cytoskeleton layer – particularly the resting length of the cytoskeleton – play key roles in specifying the degree of the membrane wrapping. We also found that the shear energy of cytoskeleton deformation diverges at the full wrapping state, suggesting the local disassembly of the cytoskeleton is required to complete the merozoite entry. Additionally, using our theoretical framework, we predict the landscape of myosin-mediated forces and the physical properties of the RBC membrane in regulating successful malaria invasion. Our findings on the crucial role of RBC membrane tension in inhibiting malaria invasion can have implications for developing novel antimalarial therapeutic or vaccine-based strategies.

**Significance:** RBC membrane tension plays an important role in regulating RBC shape and functionality. In particular, recent experimental studies have shown that elevated RBC membrane tension protects against severe malaria infection. In this study, we sought to identify how different contributions to the the effective membrane tension can contribute to this mechanically-driven protection against malaria invasion. Using a mathematical model, we derived a relationship between the effective tension of the RBC membrane – comprising a lipid bilayer and a cytoskeleton layer– and the degree of membrane wrapping during malaria invasion. Our model shows that the shear resistance of the RBC cytoskeleton plays an important role in inhibiting malaria invasion. Our findings can be generalized to the role of cell membrane mechanics in many wrapping phenomena providing insight into the crucial contributions of the host-cell membrane in protection against severe infections.

## Introduction

Malaria is one of the major infectious diseases and causes nearly half a million deaths per year worldwide [1]. Merozoites, which are protozoan parasites of the *Plasmodium* family, are small egg-shaped parasites with a diameter of 1-2 μm and are key to infecting red blood cells (RBCs) in the progression of malaria [2–4]. Merozoites invade healthy RBCs and asexually reproduce inside them; this is a critical step in the survival and reproduction of the parasites. In recent years, extensive studies have been focused on understanding the molecular and biophysical mechanisms underlying erythrocyte invasion as a route to develop novel antimalarial therapeutic or vaccine-based strategies [5–7].

At the cellular level, the invasion of erythrocytes by merozoites can be classified by the biophysical processes that include parasite binding to the RBC and subsequent membrane bending (Fig. 1A). The formation of the tight junction between the merozoite and the RBC involves low-affinity attachment of merozoite surface proteins 1 (MSP1) to RBCs, reorientation of the merozoite, and strengthening of the adhesion by binding of the erythrocyte-binding-like (EBL) or the reticulocyte binding antigen homolog (Rh) proteins and the erythrocyte membrane receptors (Fig. 1A) [8–10]. For a period of time, the penetration of the parasite into the RBC was assumed to be solely driven by the parasite actomyosin motors suggesting that the erythrocyte surface played a barrier role in the entire invasion (Fig. 1A) [11–14]. However, recent evidence has challenged this view [15–17]. Several studies have demonstrated the reorientation and the formation of host cell actin filaments dense structures at the point of entry during *Toxoplasma* and nonerythroid *Plasmodium* invasion [18–20]. Andenmatten et al. reported a degree of invasion even by knocking out the myosin and actin in *Toxoplasma*, proposing the existence of alternative invasion pathways in apicomplexan parasites [21]. Thus, all of these studies highlight the important role of the host cell during merozoite entry.

**Figure 1:**
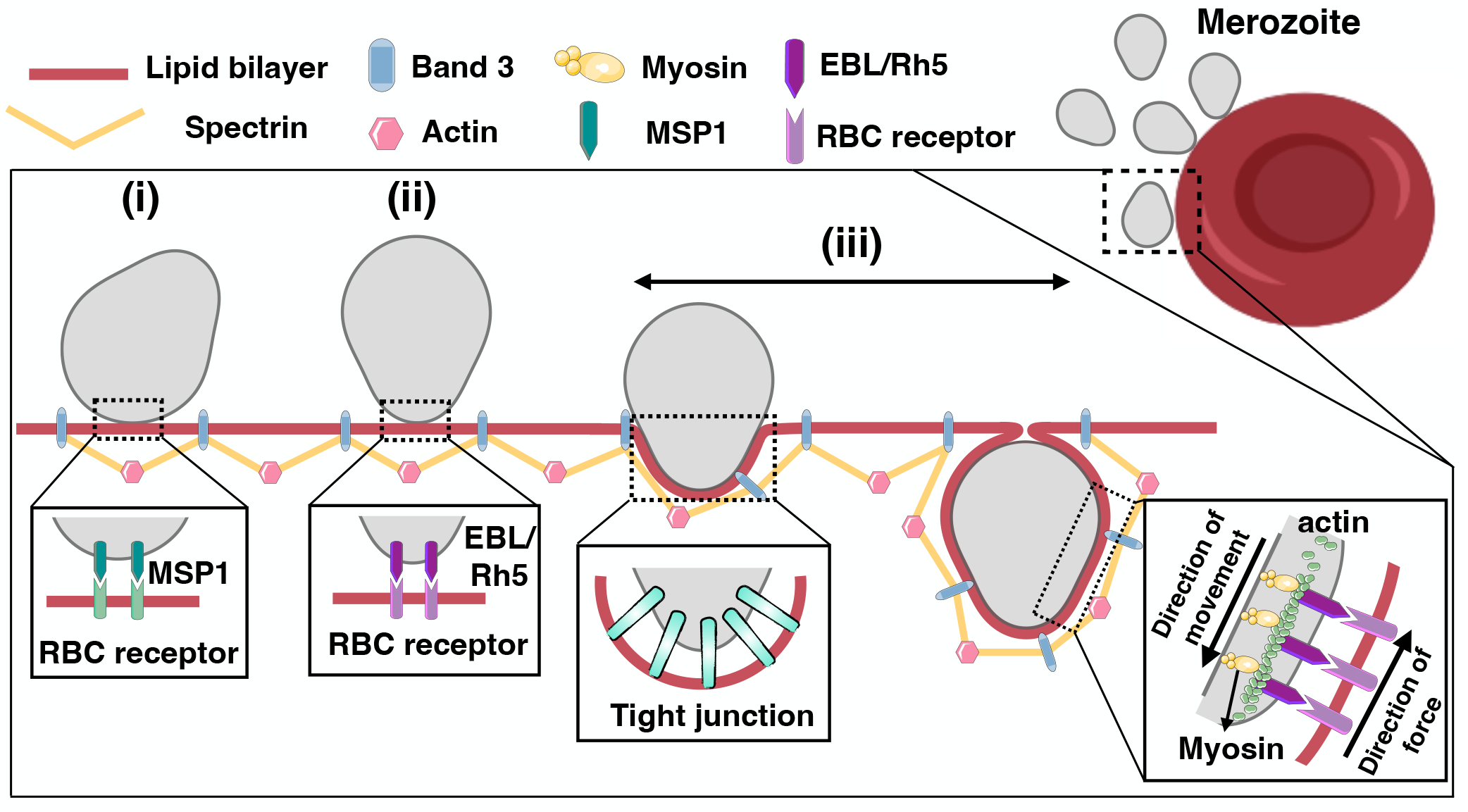
Schematic depiction of the molecular machinery of erythrocytic invasion by malaria parasites. Different stages of malaria invasion. The activity of the parasite actomyosin motors generates forces that push the merozoite into the RBC. In this study, we only focus on the erythrocyte membrane wrapping stage (iii). Msp1 is a merozoite surface protein 1, EBL is an erythrocyte-binding-like protein, and Rh is a reticulocyte binding antigen homolog protein on the malaria parasite surface.

In this study, we specifically focus on the role of the RBC membrane tension on the invasion capability of the merozoite using a continuum mechanics approach. Our study is motivated by recent evidence that the tension of the RBC membrane has been implicated as a key determinant of merozoite invasion [22]. We first identify the known main contributors to the RBC membrane tension and RBC mechanical properties from the literature and summarize their role in the success or failure of the invasion below.

### (i) Role of lipid bilayer incompressibility

The lipid bilayer in an RBC is assumed to be resistant to stretch and, therefore, areally incompressible [23, 24]. Mathematically, the tension of the bilayer is equivalent to the Lagrange multiplier used to maintain this incompressibility [25–27]. Several studies have shown that the rare Dantu variant of the glycophorin A/B receptors, which is associated with increased RBC tension, can protect against severe malaria [22, 28, 29]. Particularly, a recent experimental work by Kariuki et al. has shown the direct relationship between RBC tension and the efficiency of merozoite invasion [22]. Using the membrane flickering spectrometry technique, they demonstrated that Dantu RBCs with high tension deform less in contact with merozoites, and there is a tension threshold (< 3.8 ± 2 × 10^−7^ N/m) above which no invasion can take place [22]. Similar observations have been made about the increased susceptibility of young RBCs with lower tension to parasite *P. falciparum* invasion [30].

### (ii) Role of parasite-induced spontaneous tension

Local protein-mediated adhesion of merozoites to the surface of RBCs can induce asymmetry in lipid orientation (including lipid tilt) and distribution such as the formation of lipid rafts [31–33]. This asymmetry in the lipid membrane composition imposes a curvature on the membrane, termed spontaneous curvature [34]. The concept of spontaneous curvature has been widely used to explain different observed shapes of RBCs from stomatocytes to discocytes and echinocytes [35–39]. Additionally, Kabaso *et al*. showed that the induced local spontaneous curvature due to the spatial attachment of spectrin filaments to the inner surface of the RBC lipid bilayer is a key mechanism that drives the insideout membrane curling phenomena [40]. Despite differences in its molecular origin, spontaneous curvature is known to contribute to the membrane tension, in what is termed as spontaneous tension [41]. Dasgupta *et al*. have investigated the role of spontaneous tension in the malaria invasion by introducing an effective tension, including the contributions of both RBC membrane tension due to incompressibility and spontaneous tension [4]. They found that for high effective tension, the transition between erythrocyte wrapping states is continuous, whereas, for low effective tension, the transition is associated with an energy barrier [4].

### (iii) Role of merozoite adhesion and line tension

A critical step in malaria invasion is the adhesive interaction between the RBC membrane and the merozoite. For a full membrane-driven merozoite wrapping, the energy gained by merozoite adhesion to the RBC surface needs to overcome the energy cost due to membrane bending, membrane tension resistance, and cytoskeleton deformation [42, 43]. For instance, the absence of the Duffy antigen receptor for chemokines (DARC) on RBC surfaces significantly reduces merozoite adhesion to the membrane, which makes the Duffy-negative blood group resistant to malaria invasion [44]. Line tension at the boundary of the merozoite attachment site characterizes the discontinuity in membrane properties between the region adhering to the merozoite and the free membrane outside of the invagination domain [39]. Interfacial line tension at the merozoite boundary can create a nucleation barrier in the early stage of merozoite wrapping by increasing the energy required to form the initial membrane invagination [45]. Ignoring the effects of RBC cytoskeleton, Dasgupta et al. suggested that at low adhesion strength, interfacial forces impede the merozoite entry [4], but at high adhesion strength, these interfacial forces push the merozoite forward from partial wrapping to full wrapping [4].

### (iv) Role of the RBC cytoskeleton

The RBC cytoskeleton is a two dimensional lattice that is made of short F-actins interconnected by flexible spectrin molecules and provides support for the RBC membrane to maintain its curvature, tension, and physical properties [46–49]. The RBC cytoskeleton is a strong elastic network that restricts the deformation of the membrane and also contributes to the organization of the membrane proteins. Thus, the mechanical properties of the RBC cytoskeleton and its interaction with the lipid bilayer play important roles in malaria invasion [13, 15]. For example, several studies have identified that in an ovalocytic erythrocyte, a more rigid cytoskeleton (3-4 times higher shear modulus compared to normal cells) significantly impairs the parasite invasion process [50–52]. *In vivo*, augmented RBCs with a cytosolic polyamine (e.g., spermine) demonstrated strong resistance against malaria invasion [53]. This is because adding polyamines increases the cohesion of the cytoskeleton and, ultimately, the mechanical rigidity of the whole RBC membrane [53]. Additionally, changes induced in the cytoskeleton structure and the viscoelastic properties of the RBC membrane due to phosphorylation of transmembrane and cytoskeletal erythrocyte proteins have been shown to facilitate malaria entry [13, 15, 54].

Previous mathematical models have mainly focused on the different biomechanical aspects of the lipid bilayer in regulating malaria invasion [4, 6, 55, 56]. However, several studies have suggested that cytoskeletal remodeling and its posttranslational modifications can play crucial roles during merozoite invasion [15, 19, 50–52]. The RBC cytoskeleton is free to move in lateral directions. At the continuum level, the empirical constitutive equation based on thermodynamic invariants has been proposed by Evans and Skalak to describe the elastic energy of the cytoskeleton in the limit of small deformation [57]. This model has been extensively used, particularly in studies on red blood cell (RBC) membrane deformation in capillaries and echinocyte formation [46, 58–62]. Recently, Feng et al. proposed a microstructure-based elastic model that accounts for large cytoskeleton deformation and strain-hardening behavior at the spectrin level [63]. However, it remains unclear how the induced tension due to the coupled bilayer-cytoskeleton and the physical properties of the spectrin-actin network affect the progress of malaria invasion.

In this work, we sought to answer the following specific questions. How does RBC membrane tension including the effects of lipid bilayer incompressibility, spontaneous curvature, interfacial tension, and the cytoskeleton resistance against deformation, impact the morphological progression of parasite wrapping during malaria invasion? What are the roles of adhesion energy and interfacial line tension at the edge of merozoite in modulating these relationships? And finally, how do changes in the physical properties of RBCs alter the mechanical landscape of actomyosin forces required to complete invasion? To answer these questions, we used a general theoretical framework that incorporates the mechanics of a lipid bilayer with cytoskeleton deformation and membrane-merozoite interactions during the malaria invasion process. Our results show that the success of parasite invasion, as measured by the wrapping of the RBC membrane around the parasite, depends on the magnitude of the effective membrane tension of the RBC, which in turn depends on both the mechanics of the membrane and cytoskeleton.

## Model Development

### Assumptions

- We model the RBC membrane as a two layer manifold with one layer for the lipid bilayer and the other for the cytoskeleton (Fig. 2A). We assume that the system is at mechanical equilibrium at all times and neglect both fluctuations and inertia [64–66]. Analogous to the cup-like model [67], we assume that the free lipid bilayer and cytoskeleton outside of the parasite surface are almost flat and we only calculate the total free energy of the system on the adhered parasite area (Fig. 2B) [4, 68].
- We treat the lipid bilayer as a continuous thin elastic shell, assuming that the bilayer thickness is negligible compared to the radii of membrane curvature [34, 35]. We also assume that the lipid bilayer is incompressible and model the bending energy of the lipid bilayer using Helfrich–Canham energy, which depends on the local curvatures of the surface and bilayer properties [24, 34, 37, 69, 70].
- We treat the cytoskeleton as a triangular elastic network with two different orientations and the network bonds (mediated by spectrin) that behave as an elastic worm-like polymer (Fig. 2C) [71, 72]. This allows us to model the entropic free energy stored in the spectrin proteins using the Worm Like Chain (WLC) model [73, 74]. We also assume that the cytoskeleton convects with the bilayer, which imposes the areal incompressibility of the bilayer-cytoskeleton composite [75–79].
- We model the contact energy between the merozoite surface and erythrocyte membrane with a contact potential, assuming that adhesive strength is homogeneous on the surface of the parasite [42, 43, 80]. Additionally, to accommodate the transition from the free membrane domain to the domain where the membrane adheres to the parasite, we consider the contribution of interfacial line tension at the edge of the adhering merozoite (Fig. 2B) [4, 42, 81].
- We assume that the movement of anchored myosin motors on the polymerized actin filaments inside the parasite pushes the RBC adhered area rearwards and propels the merozoite forward into the target cell (Fig. 2B) [14]. We model the net effect of parasite motor forces as a work done on the RBC membrane and do not include the molecular details of the actomyosin assembly; these have been considered in [4, 48, 82].
- For ease of computation, we assume that a flat circular patch of a lipid bilayer and relaxed cytoskeleton deformed to fit the merozoite contour in the adhesive region (Fig. 2A) [59, 61]. We also assume that the shape of merozoite and the deformed bilayer/cytoskeleton composite are rotationally symmetric (Fig. 2B) [4].

**Figure 2:**
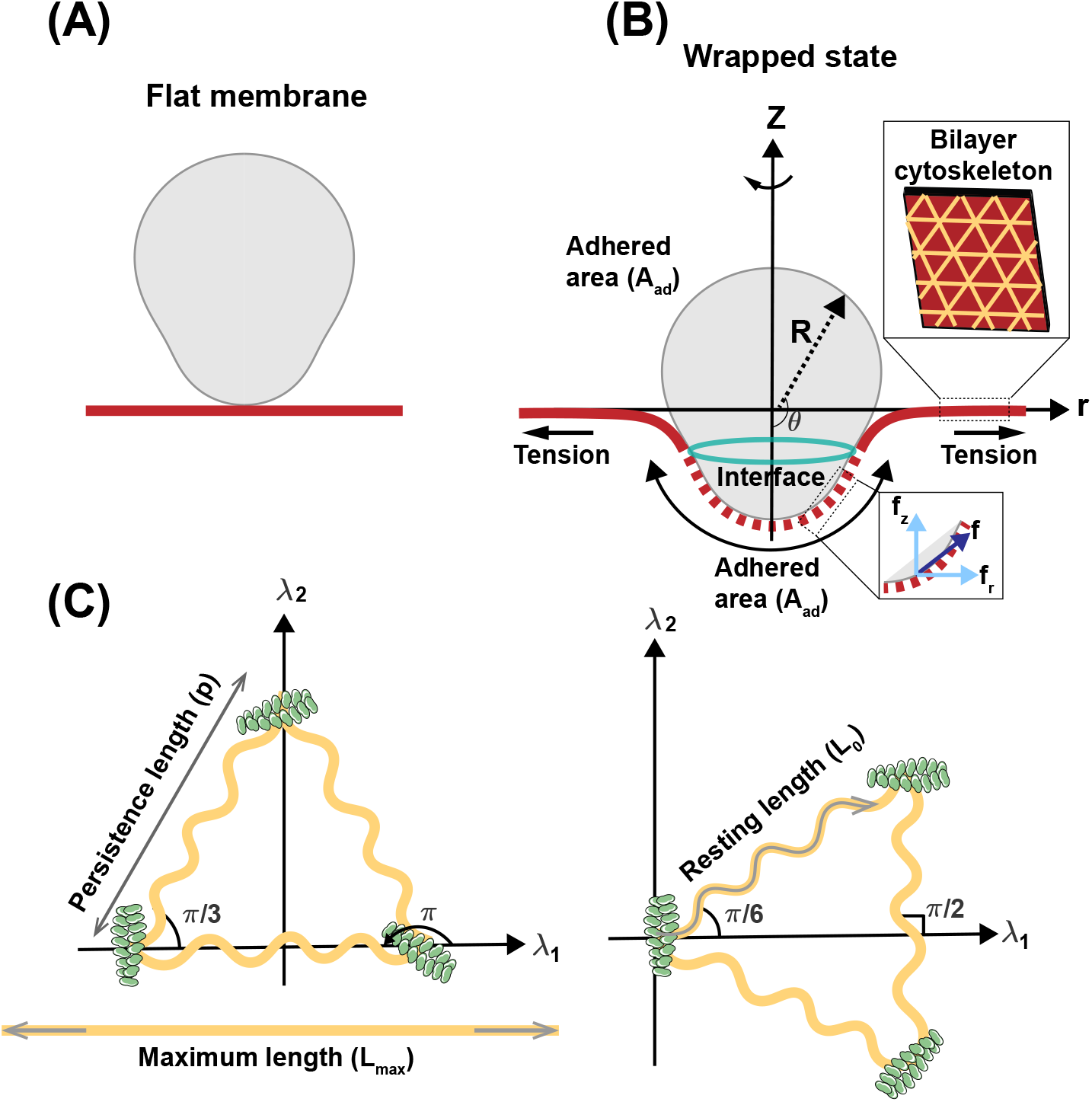
Schematic illustration of membrane wrapping and the RBC cytoskeleton. (A) Bilayer-cytoskeleton deforms from a flat circular patch to the membrane wrapping state. We model the lipid bilayer as a continuous thin elastic shell and the cytoskeleton as a hexagonal elastic network. (B) The axisymmetric parametrization of an idealized egg-shaped merozoite [4] in the wrapped state. The adhered part of the membrane is shown in dashed red, the free part of the membrane is shown in solid red, and their interface is shown by a green ring. R is the merozoite radius and *θ* is the wrapping angle. We assume that the actomyosin motors apply forces tangent to the surface (**f**) and the RBC membrane is under tension due to both lipid bilayer incompressibility and bilayer-cytoskeleton interactions. (C) The cytoskeleton network with two different orientations. *λ*_1_ and *λ*_2_ are the principal stretch directions. We model the spectrin filament as an elastic worm-like polymer with a persistence length of *p*, a maximum length of *L*_max_, and a resting length of *L*_0_ [63].

## Free energy of the system

The total energy of the system (*E*) is a sum of three terms: the energy associated with the bilayer-merozoite interactions (*E*_*b*_), the work done by the parasite actomyosin forces (*E*_*f*_), and the energy associated with the cytoskeleton deformation (*E*_*c*_)

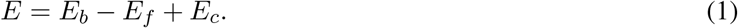

The bilayer-merozoite interactions energy includes the bending energy of the lipid bilayer, the work done against the lateral tension of the bilayer to pull excess membrane toward the parasite wrapping site, the adhesion energy due to merozoite attachment to the membrane surface, and the interfacial line tension at the edge of adhering merozoite, which is given as [4, 34, 80]

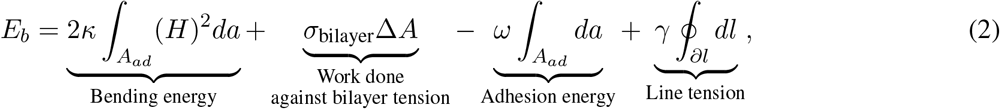

where *A*_*ad*_ is the surface area over the adhered parasite region, *H* is the membrane mean curvature, *σ*_bilayer_ is the lateral bilayer tension, Δ*A* is the excess area compared to a flat membrane, *κ* is the bending modulus of the lipid bilayer, *ω* is the adhesion energy per area, *γ* represents the strength of line tension, and the integral is over the interfacial line *dl*.

The work done on the membrane by applied forces by the parasite actomyosin motors is given by [83, 48]

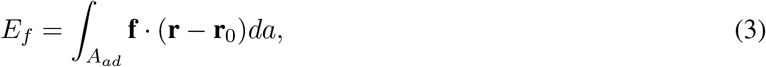

where **f** is the applied force per unit area, **r** is the position vector in the current configuration, and **r**_0_ is the position vector in the reference frame. The energy density of the membrane cytoskeleton energy including the entropic energy stored in the spectrin proteins and the steric interactions between chain elements was recently derived in [63] building on the previous models presented in [72, 84]. We expand on the details in the supplementary material and provide the free energy density per unit area here for brevity. For a 2-D triangle spectrin filament network with two different orientations, the spectrin persistence length *p*, the maximal spectrin chain length *L*_max_, the spectrin length of *L*_0_ in stress free state, the free energy density of the membrane skeleton (*W*_*c*_) can be written as (Fig. 2C) [63, 85],

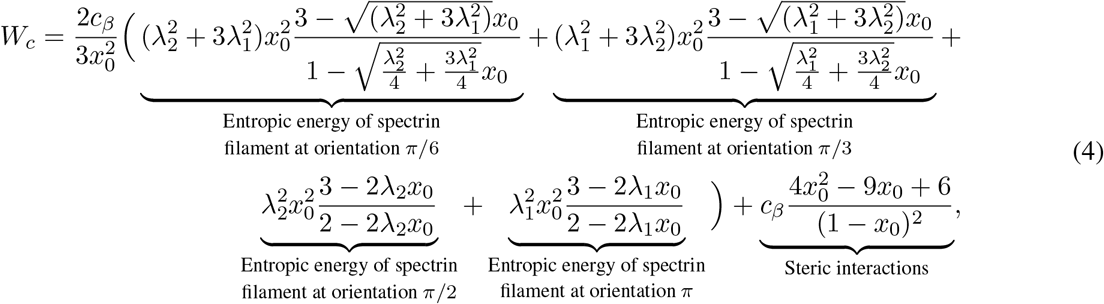

where we define *x*_0_ = *L*_0_*/L*_max_, *λ*_1,2_ are the local principal stretches, and 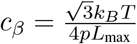(*k*_*B*_ is Boltzmann’s constant and *T* is the absolute temperature). The total energy of the skeleton can be obtained by the integral of the energy density over the adhered area to the parasite given by

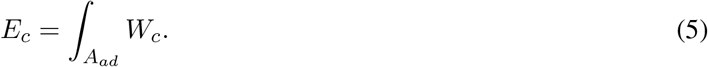

Thus, the total energy of the membrane-cytoskeleton composite is given by the sum of Eq. 2, Eq. 3, and Eq. 5. We seek to calculate the change in the energy of the system from a locally flat state to a wrapped state (Figs. 2A and B); this energy change is given by

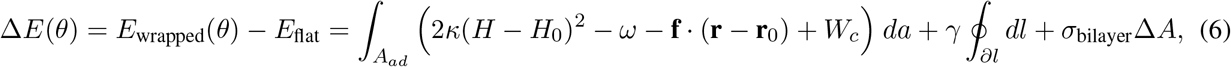

where *H*_0_ is the parasite-induced spontaneous curvature on the bilayer surface.

### Numerical implementation

A key challenge in membrane biomechanics problems is energy minimization associated with mechanical equilibrium. Traditionally, we minimize the membrane energy using the principle of virtual work to obtain the shape of the membrane in response to induced curvatures and external forces [25, 86–91]. Here, we adopt an inverse problem approach [92, 93]. Parameterizing the egg shape of the merozoite as (*X*^2^ + *Y* ^2^ + *Z*^2^) = *R*_*a*_*X*^3^ + (*R*_*a*_ − *R*_*b*_)*X*(*Y* ^2^ + *Z*^2^), with *R*_*a*_ = 1*μ*m, *R*_*b*_ = 0.7*μ*m [4], we fix the shape of the membrane adhered to the merozoite and minimize the energy (Eq. 6) associated with mechanical equilibrium to find the degree of membrane wrapping *θ*^∗^ for any given set of membrane properties. This approach has the advantage of focusing on extent of wrapping without extensive computational overhead. Biologically relevant values for the parameters that have been used in the mathematical model are listed in Table 1.

**Table 1:** Parameters used in the model.

| Parameter | Significance | Value | Ref(s) |
| --- | --- | --- | --- |
| $\kappa$ | Bending rigidity | 125 pN·nm | [22] |
| $\sigma_{\text{bilayer}}$ | Lipid bilayer tension | $10^{-4} - 1$ pN/nm | [22, 23, 94, 95, 48] |
| $\omega$ | Adhesion strength | $10^{-3} - 1$ pN/nm | [22, 96–100] |
| $\gamma$ | Interfacial force | 0 – 50 pN | [101, 102] |
| $H_0$ | Spontaneous curvature | 0 – 0.1 nm<sup>-1</sup> | |
| $L_0$ | Resting length of spectrin | 25 – 85 nm | [103–106, 85] |
| $L_{\text{max}}$ | Maximum length of spectrin | 180 – 210 nm | [107–109] |
| $p$ | Persistence length of spectrin | 10 – 25 nm | [63] |

## Results

### Successful malaria invasion is associated with an energy barrier controlled by bilayer tension

Motivated by a recent experimental observation that high RBC membrane tension can inhibit malaria invasion [22], we asked how does the interplay between the strength of merozoite adhesion and bilayer tension due to lipid incompressibility affect the degree of parasite wrapping around the RBC membrane? To answer this question, we calculated the change in the energy of bilayer/cytoskeleton composite (Eq. 6) as a function of wrapping angle (*θ*) for a fixed *σ*_bilayer_ = 0.5 pN/nm and three different magnitudes of adhesion strength (Fig. 3A). To answer this question, we calculated the change in the energy of bilayer/cytoskeleton composite (Eq. 6) as a function of wrapping angle (*θ*) for two different magnitudes of bilayer tension and adhesion strengths (Fig. 3A). Here, we set *p* = 25 nm, *L*_0_ = 35 nm, and *L*_max_ = 200 nm and there is no spontaneous curvature (*H*_0_ = 0) and no interfacial force (*γ* = 0). We find that depending on the magnitude of adhesion strength, there are three states which minimize the change in the energy; (*i*) a non-wrapped state (*θ*^∗^ = 0), (*ii*) a partially wrapped state with a small wrapping fraction (0 < *θ*^∗^ < π/4), and (*iii*) a completely wrapped state (*θ*^∗^ > *π*/2) (Fig. 3A). As can be seen in Fig. 3A, the partially wrapped state is separated from the completely wrapped state by an energy barrier. Additionally, we found that the total energy of the bilayer/cytoskeleton composite diverges toward ∞ as *θ* → *π* (Fig. 3A). This is because the first principle stretch (*λ*_1_) and the energy associated with the cytoskeleton resistance diverges for large deformations (Eq. S13). Thus, we conclude that the complete merozoite entry into the RBCs requires a local disassembly of the cytoskeleton.

**Figure 3:**
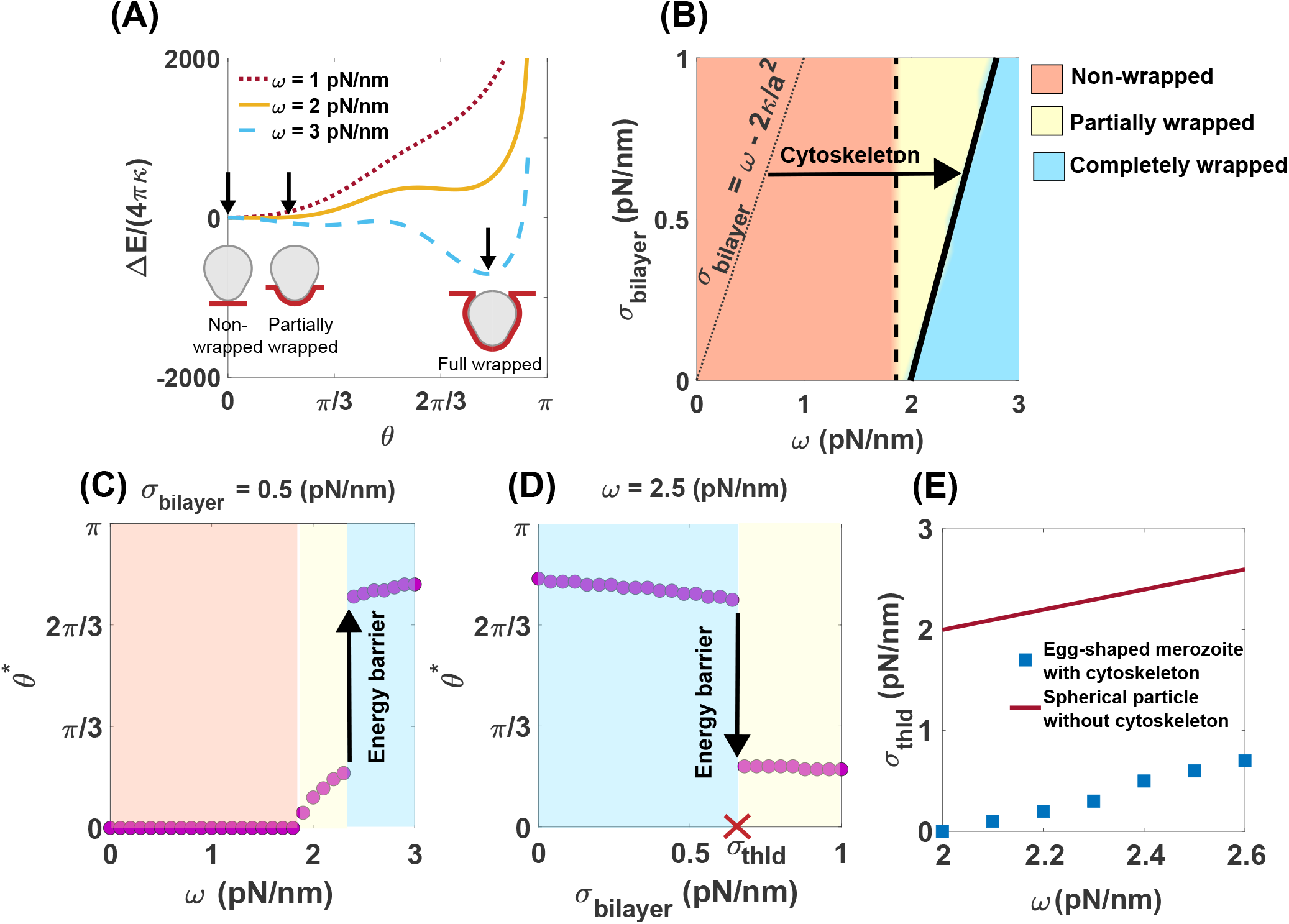
Complete merozoite wrapping by RBC membrane is associated with an energy barrier, *p* = 25 nm, *L*_0_ = 35 nm, and *L*_*max*_ = 200 nm. (A) The change in the energy of the RBC bilayer/cytoskeleton composite as a function of wrapping angle (*θ*) for a fixed *σ*_bilayer_ = 0.5 pN/nm and three different magnitudes of adhesion strength. The change in the energy is minimized either for (*i*) a non-wrapped state (*θ*^∗^ = *θ* | _minimum energy_ = 0), or (*ii*) a partially wrapped state with a small wrapping fraction (0 < *θ*^∗^ < π/4), or (*iii*) a completely wrapped state (*θ*^∗^ > *π*/2). Arrows show the location of minimum energy. (B) Merozoite wrapping phase diagram for a range of lipid bilayer tension (*σ*_bilayer_) and adhesion strength (*ω*). Orange denotes a non-wrapped state, yellow denotes a partially wrapped state, and blue are situations in which a merozoite is completely wrapped by the RBC membrane. The dashed and solid lines mark a continuous and a discontinuous transition, respectively. The dotted line represents the transition boundary below which a spherical particle is fully wrapped by the plasma membrane with no cytoskeletal effects (Eq. S17). The arrow shows the shift in the completely wrapped state toward the higher adhesion strength considering the cytoskeletal effects. (C) Wrapping angle at the minimized energy (*θ*^∗^) as a function of adhesion strength for a fixed bilayer tension (*σ*_bilayer_ = 0.5 pN/nm). An energy barrier separated a partially wrapped (failed invasion) from a completely wrapped state (successful invasion). (D) *θ*^∗^ as a function of bilayer tension for a fixed adhesion strength (*ω* = 0.2.5 pN/nm). The maximum bilayer tension required for a successful malaria invasion is marked as *σ*_thld_. (E) The bilayer tension threshold (*σ*_thld_) increases almost linearly as a function of adhesion strength.

In Fig. 3B, we plotted the merozoite wrapping phase diagram for a range of bilayer tension (*σ*_bilayer_) and adhesion strength (*ω*). The orange region denotes a non-wrapped state, the yellow region represents a partially wrapped state, and the blue region indicates a state where a merozoite is completely wrapped by the RBC membrane (Fig. 3B). We marked the continuous transition between the non-wrapped state (orange region) and the partially wrapped state (yellow region) and the discontinuous transition between the partially wrapped state and the completely wrapped state (blue region) by dashed and solid lines, respectively (Fig. 3B). In the case of a spherical particle wrapping with no cytoskeletal effects, the sphere is fully wrapped by membrane when *σ*_bilayer_ < *ω* − 2*κ/a*^2^ (shown as a dotted line in Fig. 3B). Here, *a* is the radius of the sphere with the same surface area as the egg-shaped parasite (Eq. S17) [43]. As expected, the cytoskeletal resistance against deformation shifts the transition to successful invasion toward the higher adhesion strengths (arrow in Fig. 3B), implying that larger adhesive forces are required for successful entry into an RBC. In Figs. 3C and D, we show the discontinuous transition between partially wrapped states (failed invasion) and completely wrapped states (successful invasion) with increasing the magnitude of adhesion strength and bilayer tension, respectively.

The discontinuous transition between the partially and completely wrapped states is in agreement with the proposed concept of membrane tension threshold for successful malaria invasion in a recent study by Kariuki et al.[22]. Additionally, Dasgupta et al. [4] and other studies have shown the existence of energy barriers in merozoite wrapping by RBC membrane and generally in the wrapping of nanoparticles by cellular membranes [43, 110]. We marked the maximum tension above which the invasion is impaired as *σ*_thld_ (Fig. 3D). Kariuki et al. suggested a tension threshold in order of 10^−4^ pN/nm to limit malaria invasion. However, based on our results, *σ*_thld_ is almost linearly proportional to the adhesion strength; lower *σ*_thld_ is required when the parasite adhesion strength is smaller (Fig. 3E). The linear relationship between the tension threshold and adhesion strength was expected from the analytical expression, *σ*_thld/no cytoskeleton_ ∼ *ω* (Eq. S17). However, our results show that the resistance of the cytoskeleton against deformation shifts the tension threshold to significantly lower values (Fig. 3E). Therefore, our model predicts that tension due to lipid bilayer incompressibility sets an energy barrier for a complete merozoite wrapping, while the minimum tension to impede the malaria invasion depends linearly on the strength of merozoite adhesion to the RBC membrane.

### Spontaneous tension can impede malaria invasion

It has been suggested that any asymmetry between the leaflets of the lipid bilayer or surrounding environment can induce a relatively large spontaneous tension in order of 1 pN/nm [41]. We next investigated how this induced spontaneous tension can change the efficiency of malaria invasion. Based on the analytical approximations for the membrane wrapping of a spherical particle with no cytoskeletal effects, the successful invasion occurs when *σ*_bilayer_ < *ω* − 2*κ*(1*/a* − *H*_0_)^2^ (below dotted line in Fig. 4A, Eq. S18). This suggests when the spontaneous curvature is smaller than the curvature of the merozoite (*H*_0_ < 1/*a*), the induced spontaneous tension facilitates merozoite wrapping. However, a large spontaneous curvature (*H*_0_ > 1/*a*) increases the membrane resistance to invasion. To investigate the engulfment of an egg-shaped merozoite by the RBC membrane, we set the physical properties of the membrane cytoskeleton as Fig. 3 (*p* = 25 nm, *L*_0_ = 35 nm, *L*_max_ = 200 nm) and plotted the wrapping phase diagram for a range of bilayer tension (*σ*_bilayer_) and induced spontaneous tension (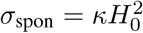, *κ* is fixed and *H*_0_ varies) at a fixed adhesion strength, *ω* = 2.5 pN/nm (Fig. 4A).

**Figure 4:**
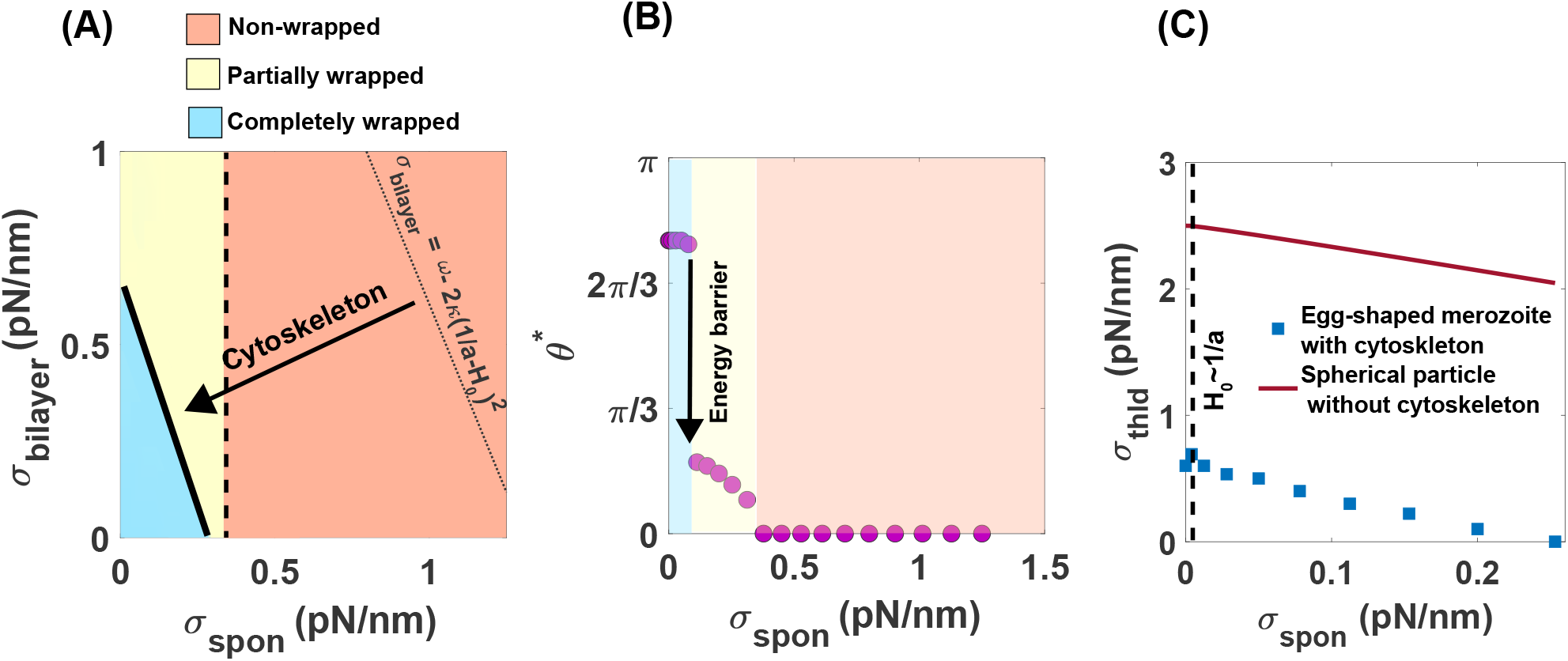
Large spontaneous tension results in malaria invasion resistance. *p* = 25 nm, *L*_0_ = 35 nm, *L*_max_ = 200 nm, and *ω* = 2.5 pN/nm. (A) Merozoite wrapping phase diagram for a range of bilayer tension (*σ*_bilayer_) and induced spontaneous tension 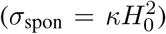 due to the local attachment of merozoites on the RBC surface. The colors indicate the same wrapping states as Fig. 3 and the solid line demonstrates the discontinuous transition between states. The dotted line represents the analytical approximation for wrapping of a spherical particle with no cytoskeletal effects (Eq. S18). The arrow shows the shift in the completely wrapped state toward the smaller spontaneous curvature due to the energy associated with the cytoskeleton deformation. (B) *θ*^∗^ as a function of induced spontaneous tension at a fixed *σ*_bilayer_ = 0.5 pN/nm. With increasing the magnitude of induced spontaneous tension, there is an energy barrier that separates the partially and completely wrapped states. (C) The nonmonotonic behavior of membrane tension threshold (*σ*_thld_) with increasing the induced spontaneous tension. When *H*_0_ < 1/*a*, *σ*_thld_ increases as the induced spontaneous tension increases, but when *H*_0_ > 1/*a*, *σ*_thld_ decreases with an increase in spontaneous tension.

Considering the energy contribution due to the elastic deformation of the membrane cytoskeleton (Eq. 5), we found that a failed malaria invasion for an egg-shaped merozoite shifts toward the smaller spontaneous tension, *σ*_spon_ < 0.5 pN/nm (arrow in Fig. 4A). To better visualize the effect of spontaneous tension on the merozoite wrapping transition, we plotted the wrapping angle (*θ*^∗^) as a function of spontaneous tension at *σ*_bilayer_ = 0.5 pN/nm (Fig. 4B). As the induced spontaneous tension increases, we observed a discontinuous transition from a completely wrapped state to a partially wrapped state and then a continuous transition from a partially wrapped state to a non-wrapped state (Fig. 4B). Based on our results, *σ*_thld_ is a nonmonotonic function of spontaneous tension; as spontaneous tension increases, *σ*_thld_ increases and then decreases again (Fig. 4C). This is consistent with the analytical approximation for wrapping of a spherical particle with no cytoskeletal effects. Thus, we predict that a large spontaneous tension can act as a protective mechanism against malaria invasion.

### Interfacial tension creates a nucleation barrier in merozoite wrapping

How do interfacial forces between the membrane at the site of invasion and the free membrane outside of adhered area influence the parasite wrapping behavior? Based on analytical approximation, the effects of interfacial forces on the wrapping process of a spherical particle with no cytoskeletal layer can be classified into three regimes (Eq. S19). (*i*) Interfacial forces create a large energy barrier (“nucleation barrier”) such that the particle does not attach to the membrane (*θ*^∗^ < 0) [42, 45, 111]. (*ii*) Interfacial forces create an energy barrier that impedes the full envelopment of a partially wrapped particle by the membrane (*θ*^∗^ < *π*/2) (Eq. S19a). (*iii*) Interfacial forces facilitate the encapsulation process when the membrane wrapping passes the equator of the sphere (*θ*^∗^ > *π*/2) (Eq. S19b) [45, 111]. Physically, the net effects of interfacial forces or line tension (*γ*) between two boundaries with different properties can be represented by an interfacial tension defined as *σ*_inter_ = *γ*^2^*/κ*. To understand how interfacial tension impacts the entry of an egg-shaped merozoite into an RBC with a cytoskeleton layer, we plotted the wrapping phase diagram for a range of bilayer tension (*σ*_bilayer_) and interfacial tension (*γ*^2^/*κ*, *κ* is fixed and *γ* varies). Here, we considered a wide range of line tension (0 < *γ* < 50 pN) to mimic line tension at lipid domain boundaries and line tension due to protein phase separation [101, 102].

We can identify three different tension regimes in Fig. 5A. At low bilayer tension (*σ*_bilayer_ < 0.5 pN/nm), independent of the magnitude of the interfacial tension, the particle is fully wrapped by the membrane. At intermediate bilayer tension (0.5 *< σ*_bilayer_ < 0.7 pN/nm), we observed a distinct contrast with the membrane wrapping of a spherical particle with no cytoskeleton layer (Eq. S19b), wherein an increase in interfacial tension results in a discontinuous transition from a completely wrapped state to a non-adhesive state (Figs. 5A and B). This discrepancy could arise from the combined effects of the merozoite’s egg-like shape and the substantial energy associated with the cytoskeleton deformation in the fully wrapped state. (Fig. 3A). In particular, for membrane wrapping of an egg-shaped merozoite with no cytoskeleton layer, we found that, depending on the combinations of the bilayer tension and adhesion strength, increasing the magnitude of interfacial tension leads to the transition of a wrapping state with *θ*^∗^ > *π*/2 to either a fully wrapped state of *θ*^∗^ = *π* (Fig. S1A) or a non-adhered state of *θ*^∗^ = 0 (Fig. S1B), which is in agreement with the previous study [4]. Finally, in the case of high bilayer tension (*σ*_bilayer_ > 0.7 pN/nm), we found that similar to the membrane wrapping of the spherical particle and the egg-shape merozoite with no cytoskeleton layer (Fig. S1C), interfacial tension creates a nucleation barrier (shown by a solid gray line in Fig. 5A) which separates a partially wrapped state from a non-adhered region (Figs. 5A and B). In Fig. 5C, we plotted *σ*_thld_ as a function of interfacial tension. Interestingly, we observed a switch-like behavior in *σ*_thld_, wherein *σ*_thld_ sharply drops from 0.6 pN/nm to 0.5 pN/nm with increasing interfacial tension (Fig. 5C). These results suggest that interfacial tension can inhibit malaria invasion by creating two energy barriers; (*1*) a nucleation barrier that impedes the merozoite attachment to the RBC membrane and (*2*) a wrapping energy barrier that hinders the full merozoite entry.

**Figure 5:**
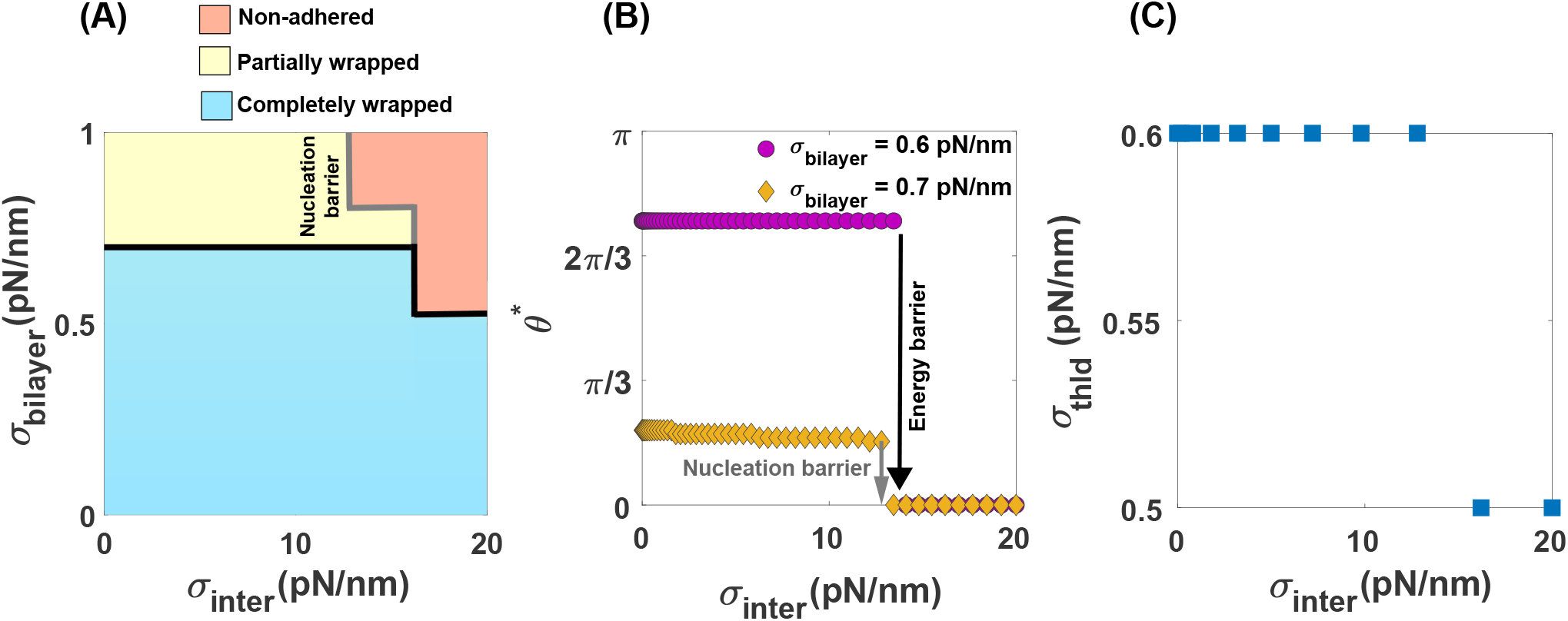
Interfacial tension plays an important role in regulating the success or failure of malaria invasion. *p* = 25 nm, *L*_0_ = 35 nm, *L*_*max*_ = 200 nm, and *ω* = 2.5 pN/nm. (A) Merozoite wrapping phase diagram for a range of bilayer tension (*σ*_bilayer_) and induced interfacial tension (*σ*_int_ = *γ*^2^*/κ*). Yellow and blue colors indicate the same wrapping states as Fig. 3, while the orange color represents a non-adhered state (*θ*^∗^ < 0). The solid lines illustrate the discontinuous transitions associated with energy barriers between different states. We marked the nucleation barrier between the partially wrapped state and the non-adhered region by a solid gray line. (B) *θ*^∗^ as a function of induced interfacial tension for two different bilayer tensions. With increasing the magnitude of induced interfacial tension, there is a discontinuous transition from completely and partially wrapped states to the non-adhered state. A switch-like behavior in the tension threshold (*σ*_thld_) with increasing the magnitude of interfacial tension.

### Physical properties of RBC cytoskeleton control the efficiency of malaria invasion

Thus far, we have fixed the properties of the membrane cytoskeleton and have focused on the role of induced tensions within the lipid bilayer in inhibiting malaria invasion. We next asked how does the tension due to the resistance of the cytoskeleton against deformation alter the degree of parasite wrapping around the RBC membrane? In this study, we assumed that the membrane cytoskeleton is areally incompressible and the spectrin network can only move in lateral directions (shear deformation). The resistance to the shear deformation is represented by the shear modulus *μ*. Initially, it has been proposed that the shear modulus of the RBC cytoskeleton is constant and is in the order of *μ* ∼ 2.5 pN*/μ*m [112–114]. However, recent studies have suggested that similar to the nonlinear response of biopolymers, the shear modulus magnitude of the spectrin network in the RBC cytoskeleton depends on the extension ratios and exhibits a strain hardening behavior in large deformations [63, 62]. For example, using a WLC model, Feng et al. derived the shear modulus of an incompressible RBC cytoskeleton as a function of principal stretches and the physical properties of the cytoskeleton (Eq. 19 in [63]).

To understand how the physical properties of the cytoskeleton affect the resistance of the RBC membrane against the malaria invasion, we fixed *H*_0_ = 0, *γ* = 0, *ω* = 2.5 pN/nm, and plotted the wrapping phase diagrams for the egg-shaped merozoite for a range of bilayer tension and natural length of spectrin (*L*_0_), the maximum length of spectrin (*L*_max_), and persistence length of spectrin (*p*) (Figs. 6A-C). We defined the average of the shear modulus of the cytoskeleton (average of *μ* in Eq. 19 in [63]) as a shear tension (*σ*_shear_). The value of the shear tension is given at the top of each panel in Figs. 6A-C. Based on our results, the merozoite wrapping process is inhibited by increasing the natural length of spectrin from *L*_0_ = 25 nm to *L*_0_ = 85 nm (Fig. 6A). This is because a larger *L*_0_ results in a higher shear tension which makes the cytoskeletal layer more rigid against deformation (Fig. 6A). In contrast to the natural length of spectrin, we found that the shear tension of the cytoskeleton decreases with an increase in the maximum or persistence length of spectrin (Figs. 6B and C). This decreases in the magnitude of shear tension facilitates the complete merozoite wrapping transition (Figs. 6B and C). It should be mentioned in all panels of Fig. 6, the transition between the completely and partially wrapped states is discontinuous (shown by a solid line), but the transition between the partial and non-wrapped states is continuous (shown by a dashed line) (see Figs. S2A-C). From these results, we can conclude that the physical properties of the cytoskeleton play key roles in specifying the magnitude of shear tension and, ultimately, the RBC resistance against malaria invasion.

**Figure 6:**
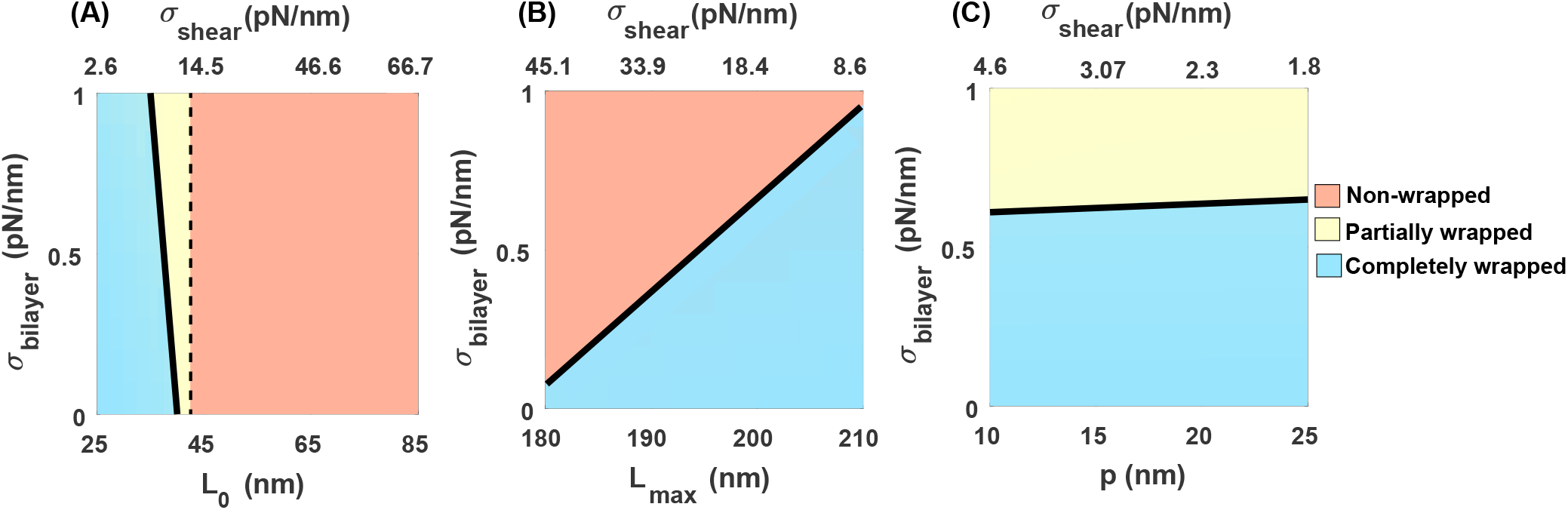
Significance of RBC cytoskeleton for inhibiting parasite invasion, *ω* = 2.5 pN/nm. Merozoite wrapping phase diagram for a range of bilayer tension and (A) the resting length of spectrin *L*_0_ (*p* = 25 nm and *L*_*max*_ = 200 nm), (B) the maximum length of spectrin *L*_max_ (*p* = 25 nm and *L*_0_ = 35 nm), and (C) the persistence length of spectrin *p* (*L*_0_ = 35 nm and *L*_*max*_ = 200 nm). In all panels, the colors show the same wrapping states as Fig. 3. The dashed and solid lines represent the continuous and discontinuous transition between the states, respectively. Shear tension (*σ*_shear_) is defined as the average shear modulus of the cytoskeleton (average of *μ* in Eq. 19 in [63]).

### Biophysical properties of RBCs alter the magnitude of actomyosin forces required for a successful malaria invasion

While the parasite’s motor forces are known as the primary driving mechanism for malaria invasion [11–14], we next asked whether the magnitude of motor-driven forces varies based on the biophysical properties of the host RBC membrane. To answer this question, we first estimated the minimum axial forces (*F*_*z*_) required for a full envelopment of a spherical particle with no cytoskeleton layer (Eq. S22). Based on our analytical approximation, *F*_*z*_ is linearly proportional to the lipid bilayer bending rigidity (*κ*), bilayer tension (*σ*), adhesion strength (*ω*), and line tension (*γ*), and it varies as a quadratic function of spontaneous curvature (*H*_0_) (Eq. S22). For example, in the case of a tensionless bilayer, with no adhesion energy, no spontaneous curvature, and no line tension, a minimum axial force of *F*_*z*_ ∼ 3 pN is required for a full wrapping of a spherical particle (Eq. S22), which is of the order of the reported actomyosin forces needed for a complete merozoite invasion by Dasgupta et al. [4]. Taking into account the energy contribution of cytoskeleton deformation, we numerically calculated the minimum axial forces required for a successful invasion by an egg-shaped merozoite, considering a range of lipid bilayer and cytoskeleton properties (Figs. 7A-F).

**Figure 7:**
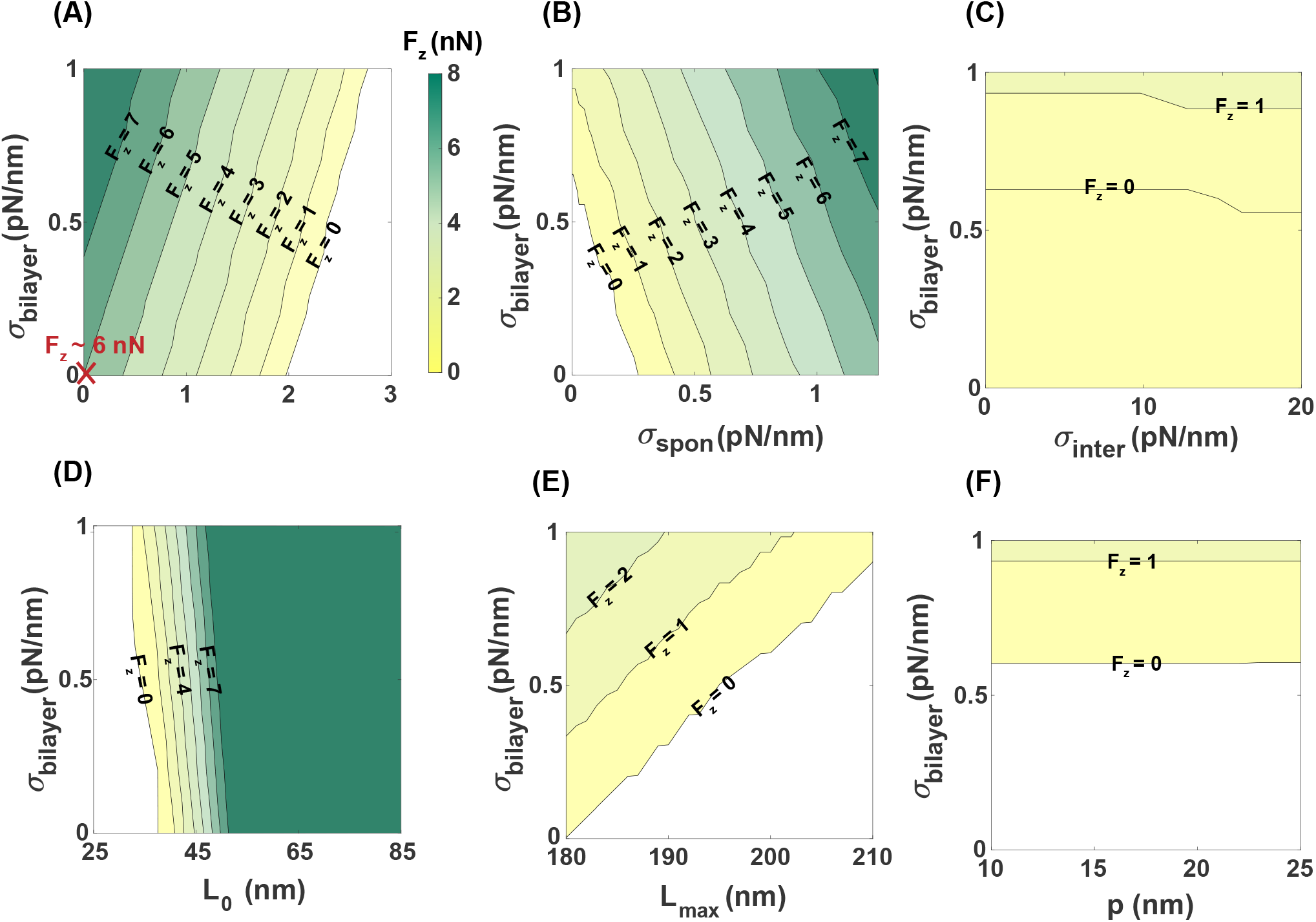
The magnitude of actomyosin forces required for a complete invasion depends on the biophysical properties of RBC membrane. Contour plots of the required actomyosin forces for a complete membrane wrapping for a range of bilayer tension and (A) the adhesion strength, (B) the spontaneous tension, (C) interfacial tension, (D) the resting length of cytoskeleton *L*_0_, (E) the maximum length of cytoskeleton *L*_*max*_, and (F) the persistence length of cytoskeleton *p*. In panels A-C, the physical properties of the cytoskeleton are set as *p* = 25 nm, *L*_0_ = 35 nm, and *L*_*max*_ = 200 nm and the results in panels B-F, are obtained for *ω* = 2.5 pN/nm. Also, in panels D-F, we set *H*_0_ = 0 and *γ* = 0 to focus on the cytoskeleton effects. The marked point **X** in panel A shows the minimum axial force (*F*_*z*_ ∼ 6 nN) required for successful invasion in a bilayer/cytoskeleton composite with a tensionless bilayer, no adhesion energy, no spontaneous curvature, and no line tension.

Consistent with the analytical approximations for a membrane wrapping of a spherical particle with no cytoskeleton layer, we observed a linear increase in the magnitude of the axial force (*F*_*z*_) with respect to bilayer tension and spontaneous tension, and a linear decrease as adhesion strength increases (Figs. 7A-B and S3A-C). Moreover, our numerical results show that *F*_*z*_ undergoes a switch-like transition from a smaller to a larger value in response to increasing interfacial tension (Figs. 7C and S3A-D). Based on our results, the magnitude of axial forces required for complete merozoite entry significantly increases when taking into account the shear energy associated with cytoskeleton deformation (Figs. 7 and S3). For example, considering the case of a bilayer/cytoskeleton composite with a tensionless bilayer, with no adhesion energy, no spontaneous curvature, and no line tension, a minimum axial force of *F*_*z*_ ∼ 6 nN is required for a successful invasion (Fig. 7A). This is almost three orders of magnitudes larger than the case with no cytoskeletal layer [4].

In Figs. 7D-F, we showed the effect of the spectrin length scales on the degree of forces required to facilitate parasite transitions to a completed invasion. As expected from Fig. 6, larger axial forces are needed for a cytoskeleton network with longer spectrin filaments in the resting condition (Fig. 7D) or shorter spectrin filaments in the maximum stretched state (Fig. 7E) and in the persistence state (Fig. 7F). Based on our results, the natural length scale of spectrin (*L*_0_) has a considerable effect –compared to the other length scales of spectrin (*L*_*max*_ and *L*_0_)– on the degree of motor forces required for a successful invasion (Figs. 7D-F). For example, for fixed bilayer properties and *L*_0_ = 25 nm, the merozoite is completely wrapped with no need for extra forces (*F*_*z*_ = 0), while at *L*_0_ = 50 nm, a minimum axial force of *F*_*z*_ ∼ 8 nN is required to push the parasite into the RBC (Fig. 7D). This is because the shear tension of the cytoskeleton strongly depends on the natural length scale of the spectrin filaments. Overall, these results indicate that the biophysical properties of the RBC lipid bilayer and cytoskeleton layer adjust the degree of the motor forces required for a complete merozoite invasion. Particularly, our mechanical model predicts that large actomyosin forces (*F*_*z*_ ∼ O (1 nN)) are needed to drive the merozoite forward into the RBC, considering the resistance of the cytoskeletal layer against deformation.

### Conclusions and Discussion

During the blood stage of malaria infection, thousands of merozoites, which are the smallest egg-shaped parasites with a typical size of 1-2 *μ*m, invade healthy RBCs and asexually reproduce inside them. The invasion process was initially assumed to be solely driven by the parasite actomyosin motor forces. However, recent experiments have shown that the biophysical properties of the RBC membrane, particularly the tension of the RBC membrane, also play an important role in controlling the malaria invasion [22, 17, 115]. The RBC membrane is a two layer manifold composed of an incompressible lipid bilayer and elastic spectrin-actin network. Previous theoretical studies for malaria invasion have not considered the active role of the RBC cytoskeleton and focused on the effect of induced tension within the lipid bilayer in regulating malaria invasion [4, 55].

Evans and Skalak initially proposed the empirical constitute elastic energy of the spectrin-actin network at the continuum level, treating the cytoskeleton as an isotropic hyperelastic material with constant shear and stretch moduli [112]. This classical model was able to explain the RBC deformability in capillaries and echinocyte formation with genetic defects [46, 58, 57, 59–62]. Subsequent extensions of this model include consideration of molecular details of the spectrin network [84, 72, 116, 117]. For example, for small deformations, Dao et al. calculated shear and area moduli of the cytoskeletal layer based on the virial stress at the spectrin level [118]. Additionally, Hendrickson et al. extended the mechanics of lipid bilayer with a conforming cytoskeletal layer [75]. They modeled the lipid bilayer as a nematic liquid crystal and assumed that the cytoskeleton is tethered to it by a so-called connector field [75]. Recently, Feng et al. derived an analytical hyperelastic constitutive model for the RBC cytoskeleton using the macroscopic behavior of spectrin filaments as a WLC [63]. Their proposed model accounts for the distribution of orientations and natural lengths of spectrin, representing the strain-hardening behavior of the RBC membrane observed in experiments [63].

In this study, we present a mathematical framework for investigating the role of tension of the RBC membrane, consisting of a lipid bilayer and a cytoskeleton layer, in governing the energy landscape of merozoite entry. Here, we modeled the lipid bilayer as an incompressible elastic shell that can bend and the cytoskeleton as an incompressible triangular elastic network (WLC model proposed by Feng el al. [63]) that can undergo shear deformation [34, 63]. Our results show that increasing the effective tension of the RBC membrane generates a wrapping energy barrier, which can hinder the merozoite invasion (Fig. 8). The effective tension of the RBC membrane, as summarized in Fig. 8 includes the RBC bilayer tension resulting from lipid incompressibility, the cytoskeleton shear tension due to its resistance to deformation, the induced spontaneous tension arising from asymmetry in the lipid distribution, and the induced interfacial tension at the lipid/protein phase separated boundaries (Fig. 8).

**Figure 8:**
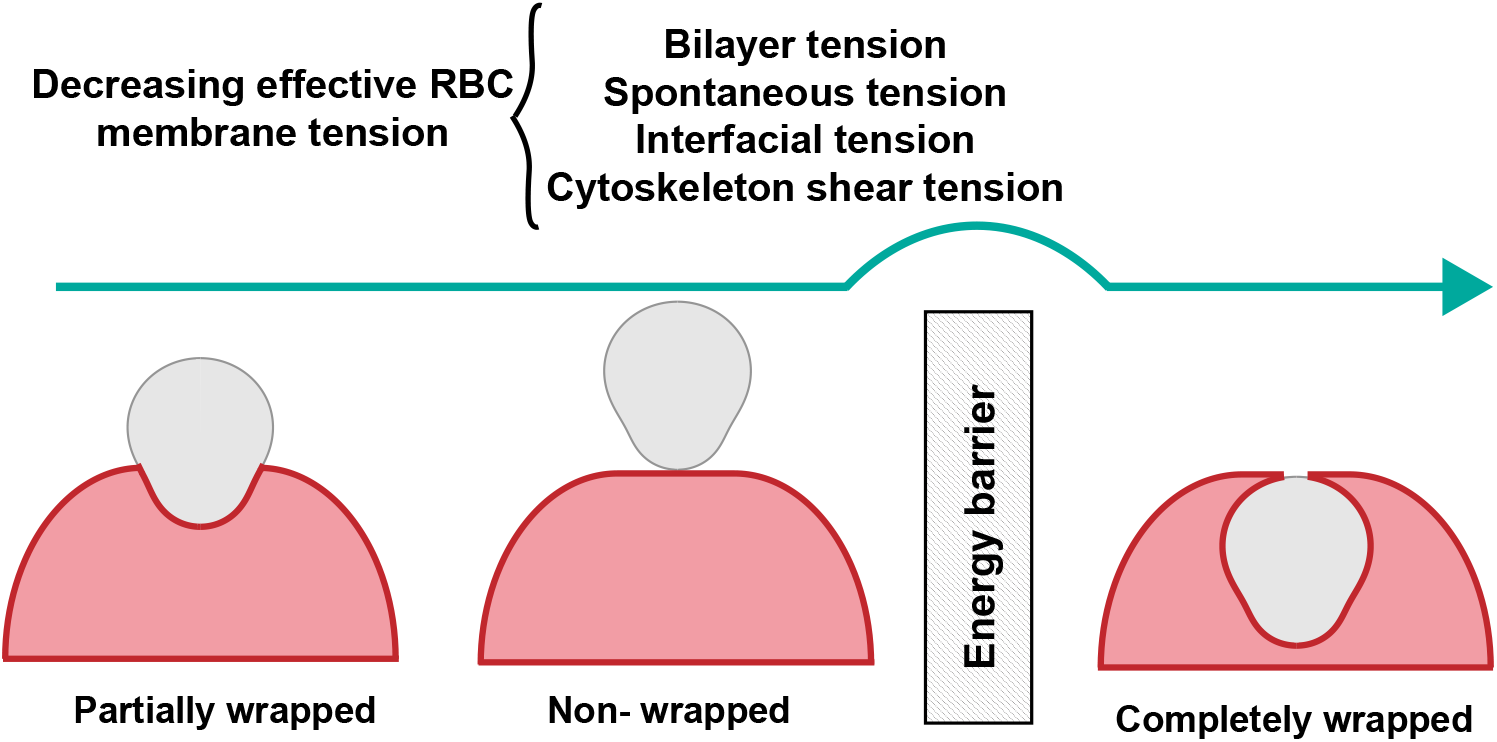
High tension of the RBC membrane, including the bilayer tension, induced spontaneous tension, interfacial tension, and the cytoskeleton-induced shear tension, acts as a protective mechanism by generating a wrapping energy barrier and inhibiting malaria invasion.

The presence of an energy barrier in the membrane wrapping process is in agreement with the concept of a membrane tension threshold required for successful malaria invasion proposed in a recent study by Kariuki et al. [22]. Based on our results, the tension threshold needed for inhibiting malaria invasion (*i*) displays an almost linear relationship with the merozoite adhesion to the RBC surface (Fig. 3), (*ii*) exhibits a non-monotonic trend with respect to spontaneous tensions, with a slight increase followed by a decrease as spontaneous tension increases (Fig. 4), and (*iii*) undergoes a sharp transition from large to small values under high interfacial tensions (Fig. 5). These results from our numerical calculations are supported by the analytical expression for membrane wrapping of a spherical particle with no cytoskeleton layer (Eqs. S17, S18, and S19).

Taking into account the energy associated with the cytoskeletal deformation in the membrane wrapping process, we found a substantial reduction in the tension threshold for impeding malaria invasion (Figs. 3 and 4). Also, our results show that the shear energy of the cytoskeleton diverges at the full wrapping state, suggesting that parasites need to utilize a mechanism to disassemble the cytoskeleton locally in order to complete the invasion process. (Figs. 3). Several experimental studies have investigated the role of the RBC cytoskeleton in mediating merozoite invasion [119–121]. In particular, in support of the local disassembly of the cytoskeleton, it has been shown that the success of malaria invasion depends on intracellular ATP hydrolysis and cytoskeleton reorganization [119–121]. We also show the correlation between the shear tension of the cytoskeleton and the three main characteristics of the spectrin network– the natural length (*L*_0_), the maximum length (*L*_max_), and the persistence length of spectrin (*p*)– in membrane wrapping progression (Fig. 6). Based on our results, increasing *L*_0_ inhibits the invasion, while larger *L*_max_ and *p* facilitate complete entry (Fig. 6). Future models should consider the molecular organization of the actin-spectrin cytoskeleton layer [49, 122, 123].

Another key aspect of malaria invasion is the role of parasite actomyosin motor forces in promoting complete entry [14, 124]. One advantage of our mathematical framework is that it can be used to estimate the minimum actomyosin motor forces required for a complete invasion while taking into account the contribution of membrane wrapping energy during merozoite entry (Fig. 7). Based on our calculations, the resistance of the cytoskeleton layer against deformation results in a significant increase (∼ three orders of magnitude) in the actomyosin force needed for complete invasion. (Fig. 7). For instance, in a case of a tensionless bilayer, with no adhesion energy, no spontaneous tension, and no interfacial tension, a minimum axial force of *F*_*z*_ ∼ 6 nN is required to deform the RBC cytoskeleton and successfully enter the host cell (Fig. 7A). Assuming that each single motor domain of the malaria parasite produces an average force of ∼ 6.5 pN [125], a minimum of ~ 920 motor domains are required to generate *F*_*z*_ ∼ 6 nN for a successful invasion. This can provide an opportunity for future studies to measure the number of active parasite motors and their relationship with the RBC membrane during malaria invasion.

The study of protection against malaria invasion has been a long-standing area of research [11–15]. Scientists have been exploring various immune responses and genetic factors to better understand and prevent this dangerous disease that kills nearly half a million people every year [126–128]. The role of RBC membrane tension in impeding malaria invasion is a new and highly intriguing concept. We believe our theoretical framework can provide insight into the mechanical aspects of the invasion, providing an opportunity for developing a novel and efficient mechanism to protect against severe malaria infection. For example, the red blood cell (RBC) membrane is characterized by a heterogeneous lipid composition, including lipid raft microdomains [129]. This heterogeneity in the lipid composition can induce a distribution of spontaneous tension across the membrane, which based on our results, could serve as a location preference for malaria invasion. Consistently, it has been shown that malarial vacuolar invaginations are enriched with the integral raft protein flotillin-1 [130]. Additionally, we have demonstrated the key contribution of the RBC cytoskeleton in inhibiting merozoite entry, which might explain the malaria protection observed in cases with mutations causing cytoskeleton rearrangements, such as individuals with sickle cell anemia and ovalocytosis[50–52, 131]. We believe these findings can be an important step toward developing efficient antimalarial therapeutic or vaccine-based strategies.

## Supplementary material for

## Specialization to axisymmetric coordinates

For computational ease, we specialize the membrane and merozoite to axisymmetric coordinates. We parameterize a surface of revolution as **r**(*s*) = *r*(*s*)**e**_*r*_ +*z*(*s*)**k**, where *s* is the arclength along the curve, *r*(*s*) is the radial distance from the axis of rotation, and *z*(*s*) is the elevation from the reference plane. Since (*dr/ds*)^2^ + (*dz/ds*)^2^ = 1, we can define *ψ* (the angle made by the tangent with respect to the horizontal) such that the normal and tangent vectors are given by **n** = − sin *ψ***e**_*r*_ + cos *ψ***k** and **a**_*s*_ = cos *ψ***e**_*r*_ + sin *ψ***k**. Following this we have *r*^∗^(*s*) = cos(*ψ*), and *z*^∗^(*s*) = sin(*ψ*) where 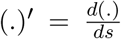. The mean curvature (*H*) can be written as 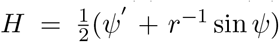. In axisymmetric coordinates, the integral over the adhered area and the integral over interfacial length in Eq. 6 simplify as

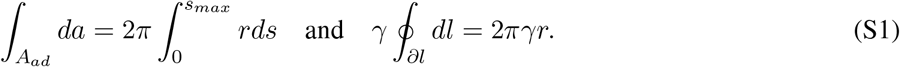

where *s*_*max*_ is the maximum membrane arclength that adheres to the merozoite. Additionally, for axisymmetric coordinates, the principal extension ratios can then be written as

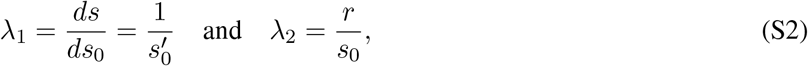

where *s*_0_ is the arclength along the undeformed shape of the axisymmetric skeleton mapping to an unknown position *s* on the deformed shape.

The egg shape of an archetypal merozoite in an axisymmetric coordinate can be parametrized as [4]

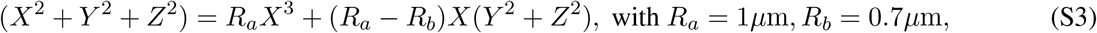

where

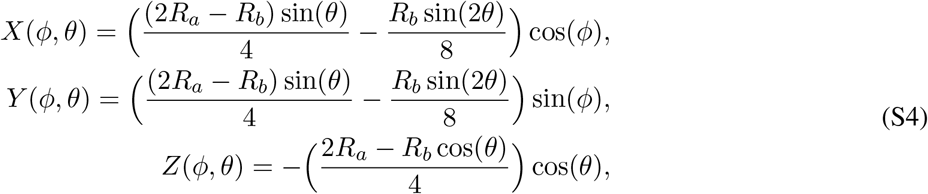

where 0 < *φ* < 2*π* and 0 < *θ* < π. Using Eq. S4, the radius of merozoite (*R*) as a function of angle *θ* can be written as

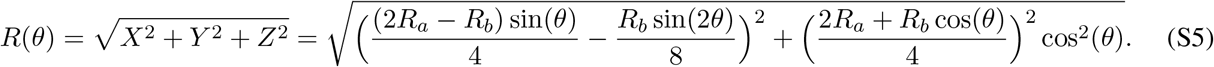

Having the radius of merozoite, we can find the radial distance from the axis of rotation (*r*) and the elevation from the reference plane (*z*) given by

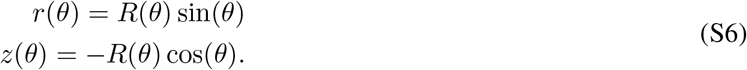

Using Eq. S6 and the definition of axisymmetric coordinates, we have

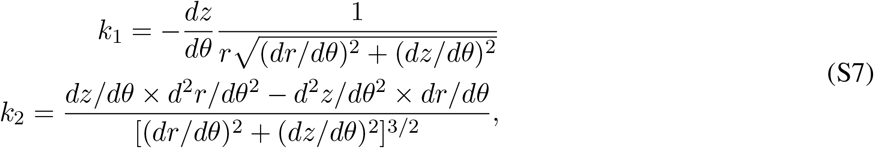

where *k*_1_ and *k*_2_ are the surface principal curvatures and 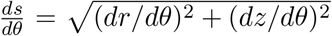. Eq. S7 allows us to find the mean curvature *H* along the merozoite surface as a function of *θ* given as

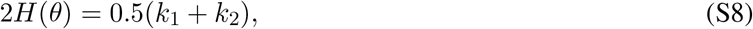

where 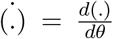. The integral over the adhered area (Eq. S1) and the extension ratios (Eq. S2) can also be calculated as a function of *θ*

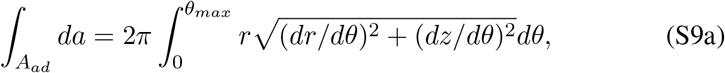

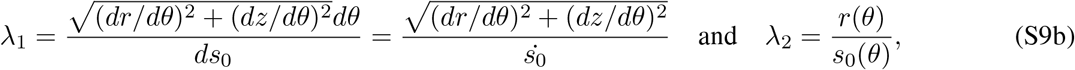

where *θ*_*max*_ is the maximum wrapping angle. Assuming that actomyosin motors apply forces tangentially along the membrane surface, in axisymmetric coordinates, the net radial force (**f**_*r*_ = *f* cos(*θ*)) is zero. Thus, only the axial component of actomyosin forces (**f**_*z*_ = *f* sin(*θ*)) pushes the merozoite forward and the work on the membrane (Eq. 3) is simplified as

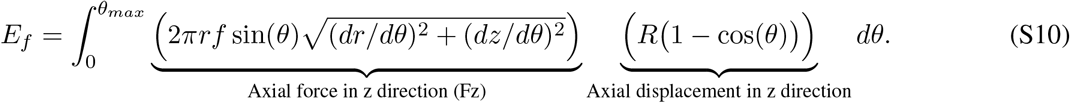

Using Eqs. S1, S9a, and S10, the change in the energy of bilayer/cytoskeleton due to the adhesion of merozoite and deformation bilayer/cytoskeleton (Eq. 6) can be written as a function *θ*

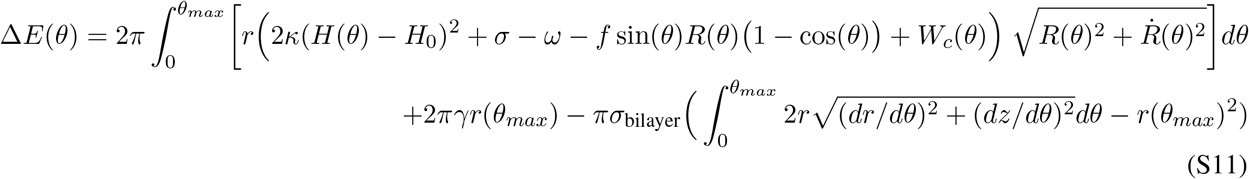

Here, we assumed that the force density applied by the actomyosin motor (*f*) is constant all along the area of adhered merozoite.

## Incompressible bilayer and cytoskeleton

Let us assume that a flat circular patch of a lipid bilayer and relaxed cytoskeleton with radius (*s*_0_) deformed to fit the merozoite contour in the adhesive region. Thus, for an incompressible bilayer/cytoskeleton, the area conservation can be written as

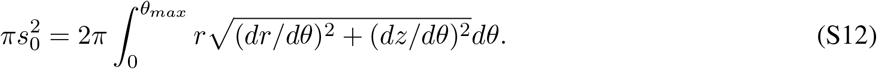

Eq. S12 allows us to find *s*_0_ and calculate the extension ratios using Eq. S9b, which simplifies as

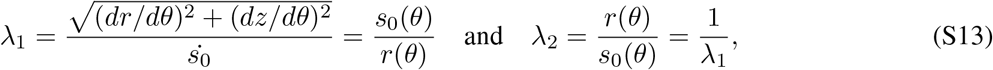

which is consistent with zero local area strain (*α* = *λ*_1_*λ*_2_ − 1 = 0) for an incompressible bilayer/cytoskeleton [112]. Additionally, for an incompressible cytoskeleton, the shear modulus *μ* is simplified as [63]

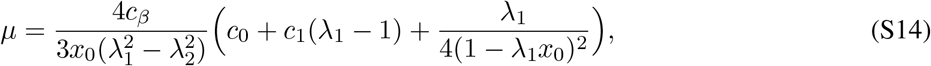

where 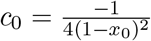 and 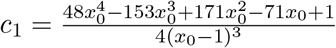

## Numerical implementation

For an egg shape merozoite parametrized by Eq. S3, we numerically calculate the change in the energy of the bilayer/cytoskeleton as a function of wrapping angle *θ* (Eq. S11). Then, for any given set of constant parameters, we find an angle *θ*^∗^ at which the invasion state becomes an energy minimum.

## Analytical approximations

In this section, we explore the analytical solution for the minimum energy state, ignoring the effects of membrane cytoskeleton energy and modeling the merozoite as a spherical particle with radius *a*. In this condition, the change in the energy of the system (Eq. S11) can be written as

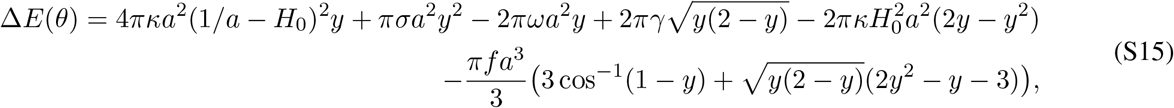

where *y* = 1 − cos(*θ*). By taking 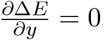, we have

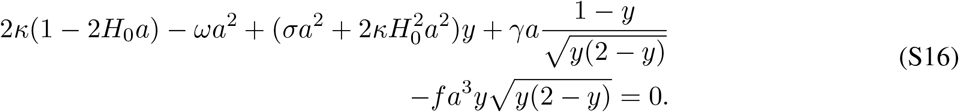

Considering our definition for a completely wrapped state (*θ*^∗^ *> π/*2), we can find the transition condition to the completely wrapped state by setting *y* = 1 in Eq. S16. Below, we simplified Eq. S16 for different conditions.

### Case 1: Relationship between lipid bilayer tension and adhesion strength

Considering the condition that *γ* = 0, *f* = 0, and *H*_0_ = 0, Eq. S16 simplifies as

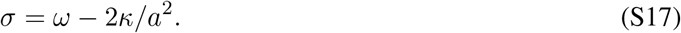

Eq. S17 suggests that the particle can get fully wrapped with increasing adhesion strength.

### Case 2: Relationship between lipid bilayer tension and spontaneous curvature

Considering the condition that *γ* = 0 and *f* = 0, Eq. S16 gives

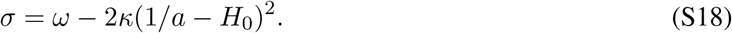

Based on Eq. S18, when the induced spontaneous curvature is smaller than the curvature of the particle (*H*_0_ < 1*/a*), the induced spontaneous curvature assists the progress of complete particle wrapping. However, larger spontaneous curvatures (*H*_0_ > 1/*a*) impede the complete wrapping transition.

### Case 3: Relationship between lipid bilayer tension and interfacial forces

As can be seen, Eq. S16 has a symmetric barrier at *θ* = *π/*2 (the line tension term vanishes for *y* = 1). Thus, to find the analytical approximation, we set *y* = 1 ± *ϵ*, where *ϵ* is a small number, and expanded the Eq. S16 until the first order for the case that *f* = 0 and *H*_0_ = 0

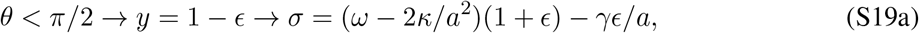

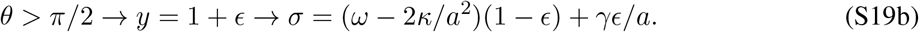

Based on Eqs. S19, in the first half of wrapping (*θ* < π/2), a line tension prevents the complete membrane wrapping process. However, once the equator is passed (*θ > π/*2), a line tension accommodates the particle encapsulation. It should be mentioned that with no line tension (*γ* = 0), a non-wrapped state (*θ*^∗^ = 0) is always a local minimum of Δ*ϵ*(*θ*). However, the line tension and actomyosin force energy terms scale as 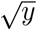 and their derivatives diverge at *θ* = 0. This means that a line tension can create an energy barrier with no minimum energy state between 0 ≤ *θ* ≤ *π* in which the particle even does not adhere to the membrane.

### Case 4: Motor forces required for a complete wrapping as a function of membrane physical properties

To calculate the minimum motor forces that are required for a complete particle wrapping (based on our definition *θ*^∗^ > π/2), we substitute *y* = 1 + *ϵ* in Eq. S16 and find the force density (*f*) as

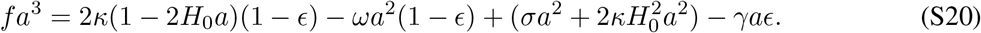

The total force in the *z* direction (*F*_*z*_) is obtained as

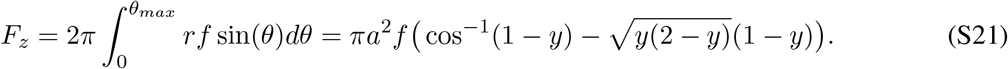

Substituting Eq. S20 into Eq. S21 for a complete wrapping condition (*y* = 1 + *ϵ*), we have

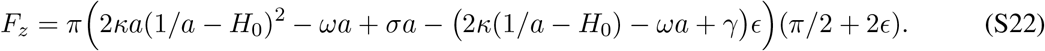

Based on Eq. S22, for a tensionless membrane (*σ* = 0), with no adhesion energy (*ω* = 0), no line tension (*γ* = 0), and no spontaneous curvature (*H*_0_ = 0), a minimum force of *F*_*z*_ = 3 pN is required for a complete wrapping of a spherical particle. This is consistent with the calculated magnitude of actomyosin forces required for a merozoite invasion by Dasgupta et al. [4]. It should be mentioned that Eq. S22 is derived for the minimum axial force needed for a complete invasion. This means if the right hand side of Eq. S22 becomes negative, the physical forces are enough to push the merozoite into the RBC and thus *F*_*z*_ = 0.

## Supplementary figures

**Figure S1:**
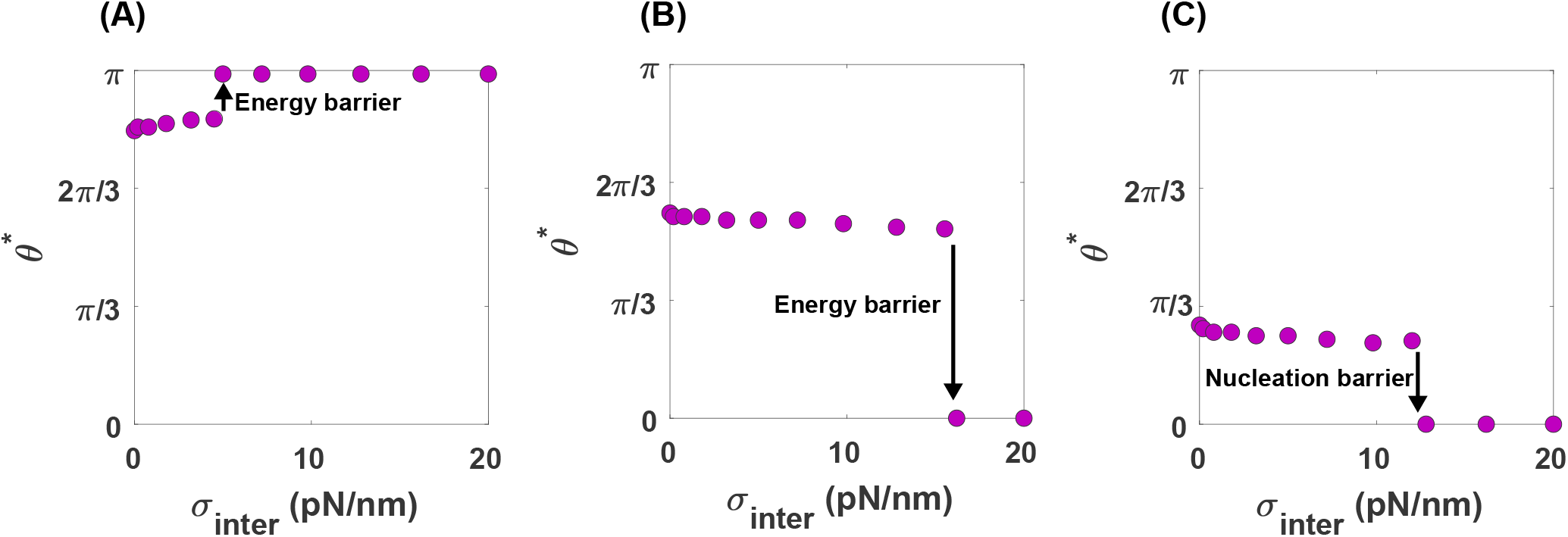
*θ*^∗^ as a function of interfacial tension for wrapping of an egg-shaped merozoite without the cytoskeleton layer. (A) A discontinuous transition from *θ*^∗^ ∼ 5*π/*6 to a full wrapped state (*θ*^∗^ = *π*) with an increase in the magnitude of interfacial tension, *ω* = 1 pN/nm and *σ*_bilayer_ = 0.6 pN/nm. (B) A discontinuous transition from *θ*^∗^ ∼ 5*π/*9 to a non-adhered state with an increase in the magnitude of interfacial tension, *ω* = 0.4 pN/nm and *σ*_bilayer_ = 0.6 pN/nm. (C) A discontinuous transition from a partially wrapped state to a non-adhered state with an increase in the magnitude of interfacial tension, *ω* = 0.5 pN/nm and *σ*_bilayer_ = 1 pN/nm.

**Figure S2:**
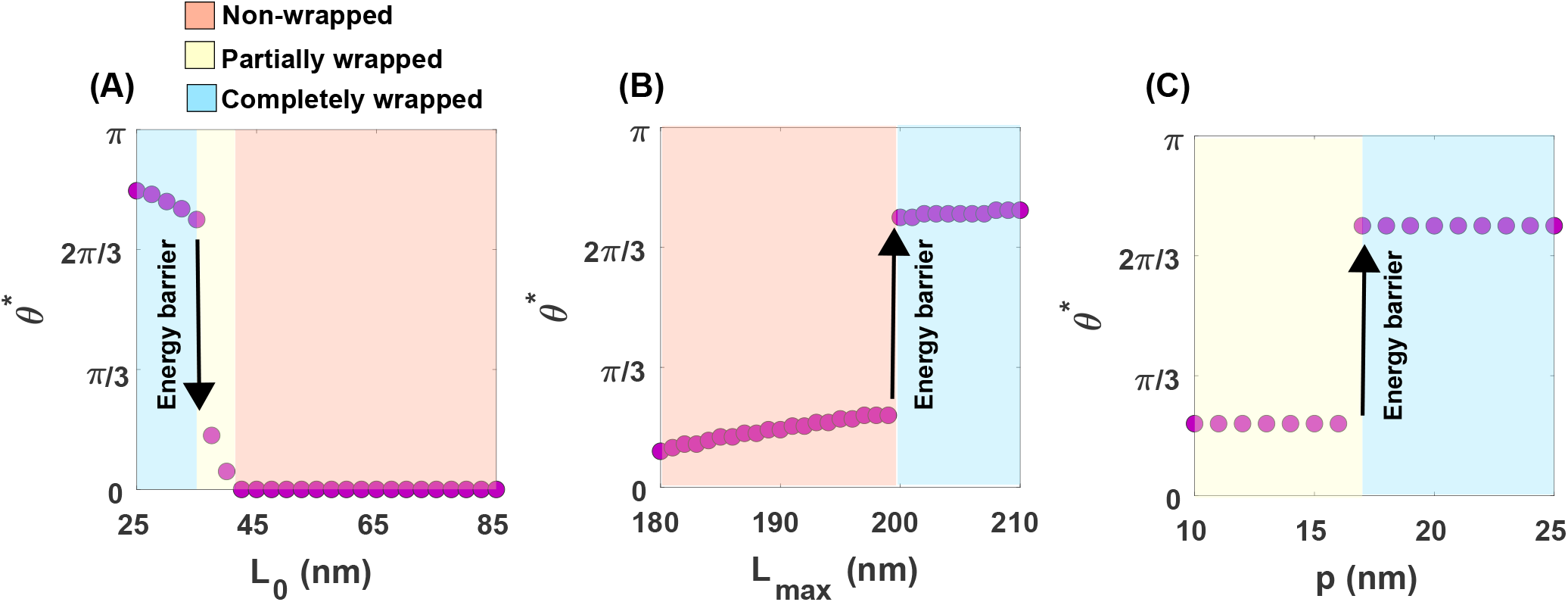
The effects of physical properties of the cytoskeleton on the efficiency of malaria invasion, *σ*_*bilayer*_ = 0.63 pN/nm. (A) A discontinuous transition from a completely to a partially wrapped state followed by a continuous transition from a partially to a non-wrapped wrapped state with increasing *L*_0_ from 25 nm to 85 nm. *σ* = 0.1 pN/nm, *ω* = 0.8 pN/nm, *p* = 25 nm, and *L*_*max*_ = 200 nm. (B) A continuous transition from a partially to a completely wrapped state followed by a discontinuous transition from a partially to a completely wrapped state with increasing *L*_*max*_ from 180 nm to 210 nm. *σ* = 0.1 pN/nm, *ω* = 0.8 pN/nm, *p* = 25 nm, and *L*_0_ = 35 nm. (C) A discontinuous transition from a partially wrapped to a completely wrapped state with increasing the persistence length of spectrin *p. σ* = 0.1 pN/nm, *ω* = 0.8 pN/nm, *L*_0_ = 35 nm, and *L*_*max*_ = 200 nm.

**Figure S3:**
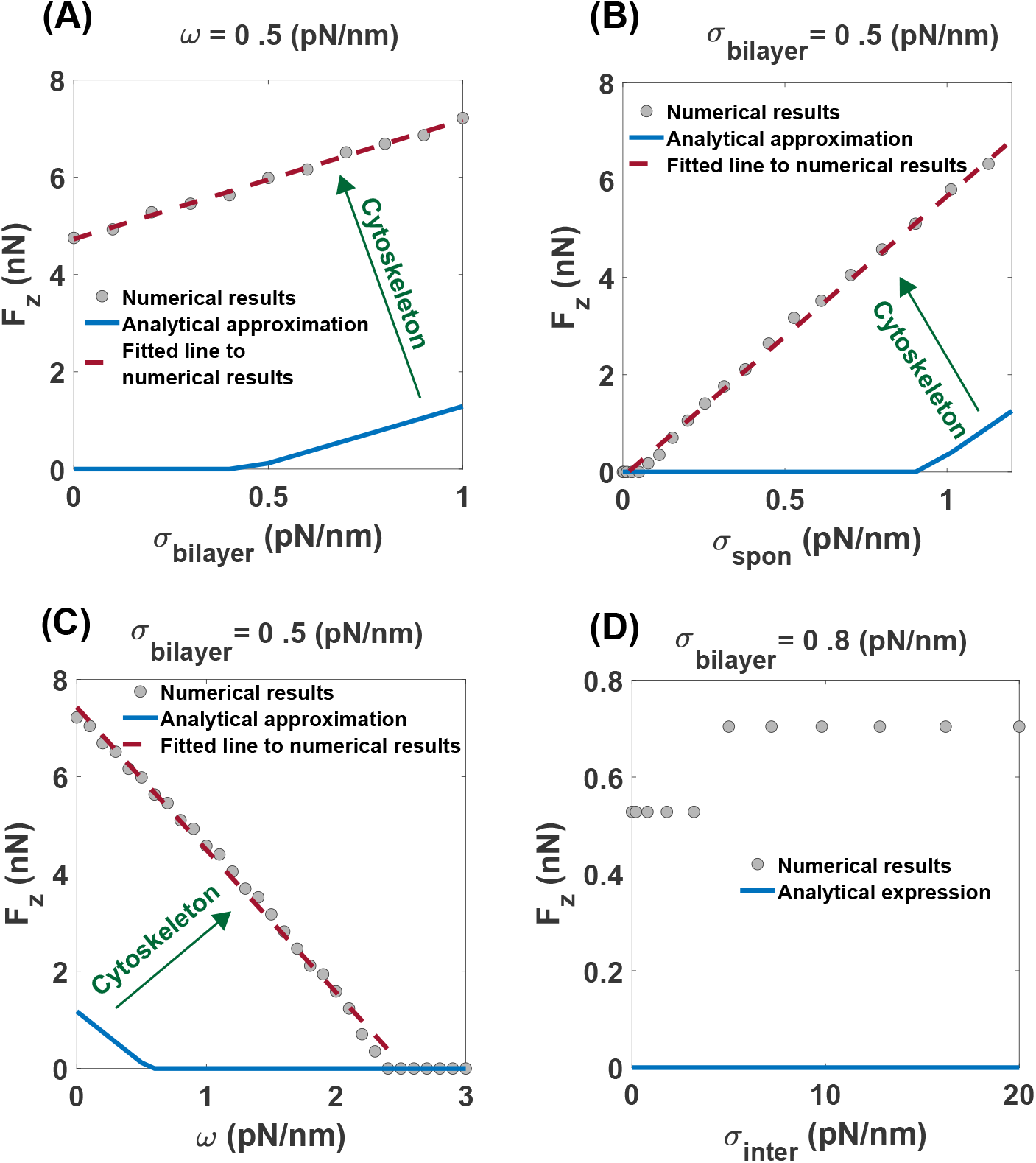
Minimum axial force (*F*_*z*_) required for a complete merozoite entry as a function of (A) bilayer tension, (B) spontaneous tension, (C) adhesion strength, and (D) interfacial tension. *p* = 25 nm, *L*_0_ = 35 nm, and *L*_*max*_ = 200 nm. The gray circles show the results that we obtained from the energy minimization (Eq. S11). The dotted line represents the fitted curves and the solid blue line indicates the analytical approximation for the motor-driven force (Eq. S22). The green arrow demonstrates the increase in the magnitude of the axial force compared to the analytical approximations because of the cytoskeleton resistance against deformation. (A) *F*_*z*_ increases as a linear function of bilayer tension. The dashed line shows the linear dependence on the bilayer tension by fitting to a line (A*σ*_bilayer_+B), where A = 2.48 and B = 4.73 with *R*^2^ = 0.99. (B) *F*_*z*_ varies as a linear function of spontaneous tension. The dashed line shows a linear dependence on the spontaneous tension by fitting to a line (A*σ*_spon_+B), where A = 5.7, B =-0.09 with *R*^2^ = 0.99. (C) *F*_*z*_ decreases as a linear function of adhesion strength. The dashed line shows the linear dependence on the adhesion strength by fitting to the line (A*ω*+B), where A = -2.93 and B = 7.4 with *R*^2^ = 0.99 (D) Switch-like increases in axial force from *F*_*z*_ = 0.52 nN to *F*_*z*_ = 0.7 nN with increasing the magnitude of interfacial tension.

## References

[1] W. H. Organization et al., “World malaria report 2020: 20 years of global progress and challenges,” 2020.

[2] E. Hanssen, C. Dekiwadia, D. T. Riglar, M. Rug, L. Lemgruber, A. F. Cowman, M. Cyrklaff, M. Kudryashev, F. Frischknecht, J. Baum, et al., “Electron tomography of p lasmodium falciparum merozoites reveals core cellular events that underpin erythrocyte invasion,” Cellular microbiology, vol. 15, no. 9, pp. 1457–1472, 2013.

[3] L. Bannister, G. Mitchell, G. Butcher, E. Dennis, and S. Cohen, “Structure and development of the surface coat of erythrocytic merozoites of plasmodium knowlesi,” Cell and tissue research, vol. 245, no. 2, pp. 281–290, 1986.

[4] S. Dasgupta, T. Auth, N. S. Gov, T. J. Satchwell, E. Hanssen, E. S. Zuccala, D. T. Riglar, A. M. Toye, T. Betz, J. Baum, et al., “Membrane-wrapping contributions to malaria parasite invasion of the human erythrocyte,” Biophysical journal, vol. 107, no. 1, pp. 43–54, 2014.

[5] A. F. Cowman, D. Berry, and J. Baum, “The cellular and molecular basis for malaria parasite invasion of the human red blood cell,” Journal of cell Biology, vol. 198, no. 6, pp. 961–971, 2012.

[6] S. Hillringhaus, A. K. Dasanna, G. Gompper, and D. A. Fedosov, “Stochastic bond dynamics facilitates alignment of malaria parasite at erythrocyte membrane upon invasion,” Elife, vol. 9, p. e56500, 2020.

[7] D. Nureye and S. Assefa, “Old and recent advances in life cycle, pathogenesis, diagnosis, prevention, and treatment of malaria including perspectives in ethiopia,” The Scientific World Journal, vol. 2020, 2020.

[8] A. F. Cowman, C. J. Tonkin, W.-H. Tham, and M. T. Duraisingh, “The molecular basis of erythrocyte invasion by malaria parasites,” Cell host & microbe, vol. 22, no. 2, pp. 232–245, 2017.

[9] P. R. Gilson and B. S. Crabb, “Morphology and kinetics of the three distinct phases of red blood cell invasion by plasmodium falciparum merozoites,” International journal for parasitology, vol. 39, no. 1, pp. 91–96, 2009.

[10] D. T. Riglar, D. Richard, D. W. Wilson, M. J. Boyle, C. Dekiwadia, L. Turnbull, F. Angrisano, D. S. Marapana, K. L. Rogers, C. B. Whitchurch, et al., “Super-resolution dissection of coordinated events during malaria parasite invasion of the human erythrocyte,” Cell host & microbe, vol. 9, no. 1, pp. 9–20, 2011.

[11] J. M. Dobrowolski and L. D. Sibley, “Toxoplasma invasion of mammalian cells is powered by the actin cytoskeleton of the parasite,” Cell, vol. 84, no. 6, pp. 933–939, 1996.

[12] J. H. Morisaki, J. E. Heuser, and L. D. Sibley, “Invasion of toxoplasma gondii occurs by active penetration of the host cell,” Journal of cell science, vol. 108, no. 6, pp. 2457–2464, 1995.

[13] J. D. Warncke and H.-P. Beck, “Host cytoskeleton remodeling throughout the blood stages of plasmodium falciparum,” Microbiology and Molecular Biology Reviews, vol. 83, no. 4, pp. e00013–19, 2019.

[14] J. Baum, A. T. Papenfuss, B. Baum, T. P. Speed, and A. F. Cowman, “Regulation of apicomplexan actin-based motility,” Nature Reviews Microbiology, vol. 4, no. 8, pp. 621–628, 2006.

[15] E. S. Zuccala and J. Baum, “Cytoskeletal and membrane remodelling during malaria parasite invasion of the human erythrocyte,” British journal of haematology, vol. 154, no. 6, pp. 680–689, 2011.

[16] M. Koch and J. Baum, “The mechanics of malaria parasite invasion of the human erythrocyte–towards a reassessment of the host cell contribution,” Cellular microbiology, vol. 18, no. 3, pp. 319–329, 2016.

[17] P. V. Groomes, U. Kanjee, and M. T. Duraisingh, “Rbc membrane biomechanics and plasmodium falciparum invasion: Probing beyond ligand–receptor interactions,” Trends in Parasitology, 2022.

[18] V. Delorme-Walker, M. Abrivard, V. Lagal, K. Anderson, A. Perazzi, V. Gonzalez, C. Page, J. Chauvet, W. Ochoa, N. Volkmann, et al., “Toxofilin upregulates the host cortical actin cytoskeleton dynamics, facilitating toxoplasma invasion,” Journal of cell science, vol. 125, no. 18, pp. 4333–4342, 2012.

[19] V. Gonzalez, A. Combe, V. David, N. A. Malmquist, V. Delorme, C. Leroy, S. Blazquez, R. Ménard, and Tardieux, “Host cell entry by apicomplexa parasites requires actin polymerization in the host cell,” Cell host & microbe, vol. 5, no. 3, pp. 259–272, 2009.

[20] M. Bichet, C. Joly, A. H. Henni, T. Guilbert, M. Xémard, V. Tafani, V. Lagal, G. Charras, and I. Tardieux, “The toxoplasma-host cell junction is anchored to the cell cortex to sustain parasite invasive force,” BMC biology, vol. 12, no. 1, pp. 1–21, 2014.

[21] N. Andenmatten, S. Egarter, A. J. Jackson, N. Jullien, J.-P. Herman, and M. Meissner, “Conditional genome engineering in toxoplasma gondii uncovers alternative invasion mechanisms,” Nature methods, vol. 10, no. 2, pp. 125–127, 2013.

[22] S. N. Kariuki, A. Marin-Menendez, V. Introini, B. J. Ravenhill, Y.-C. Lin, A. Macharia, J. Makale, M. Tendwa, W. Nyamu, J. Kotar, et al., “Red blood cell tension protects against severe malaria in the dantu blood group,” Nature, pp. 1–5, 2020.

[23] R. P. Rand and A. Burton, “Mechanical properties of the red cell membrane: I. membrane stiffness and intracellular pressure,” Biophysical journal, vol. 4, no. 2, pp. 115–135, 1964.

[24] W. Rawicz, K. Olbrich, T. McIntosh, D. Needham, and E. Evans, “Effect of chain length and unsaturation on elasticity of lipid bilayers,” Biophysical journal, vol. 79, no. 1, pp. 328–339, 2000.

[25] D. Steigmann, “Fluid films with curvature elasticity,” Archive for Rational Mechanics and Analysis, vol. 150, no. 2, pp. 127–152, 1999.

[26] M. Deserno, “Fluid lipid membranes: From differential geometry to curvature stresses,” Chemistry and physics of lipids, vol. 185, pp. 11–45, 2015.

[27] P. Rangamani, K. K. Mandadap, and G. Oster, “Protein-induced membrane curvature alters local membrane tension,” Biophysical journal, vol. 107, no. 3, pp. 751–762, 2014.

[28] S. Field, E. Hempelmann, B. Mendelow, and A. Fleming, “Glycophorin variants and plasmodium falciparum: protective effect of the dantu phenotype in vitro,” Human genetics, vol. 93, no. 2, pp. 148–150, 1994.

[29] E. M. Leffler, G. Band, G. B. Busby, K. Kivinen, Q. S. Le, G. M. Clarke, K. A. Bojang, D. J. Conway, M. Jallow, F. Sisay-Joof, et al., “Resistance to malaria through structural variation of red blood cell invasion receptors,” Science, vol. 356, no. 6343, 2017.

[30] G. Pasvol, D. Weatherall, and R. Wilson, “The increased susceptibility of young red cells to invasion by the malarial parasite plasmodium falciparum,” British journal of haematology, vol. 45, no. 2, pp. 285–295, 1980.

[31] A. Tian and T. Baumgart, “Sorting of lipids and proteins in membrane curvature gradients,” Biophysical journal, vol. 96, no. 7, pp. 2676–2688, 2009.

[32] L. V. Schäfer and S. J. Marrink, “Partitioning of lipids at domain boundaries in model membranes,” Biophysical journal, vol. 99, no. 12, pp. L91–L93, 2010.

[33] P. Rangamani and D. Steigmann, “Variable tilt on lipid membranes,” Proceedings of the Royal Society A: Mathematical, Physical and Engineering Sciences, vol. 470, no. 2172, p. 20140463, 2014.

[34] W. Helfrich, “Elastic properties of lipid bilayers: theory and possible experiments,” Zeitschrift fur Naturforschung C, vol. 28, no. 11–12, pp. 693–703, 1973.

[35] H. Deuling and W. Helfrich, “Red blood cell shapes as explained on the basis of curvature elasticity,” Biophysical journal, vol. 16, no. 8, pp. 861–868, 1976.

[36] G. L. Hw, M. Wortis, and R. Mukhopadhyay, “Stomatocyte–discocyte–echinocyte sequence of the human red blood cell: Evidence for the bilayer–couple hypothesis from membrane mechanics,” Proceedings of the National Academy of Sciences, vol. 99, no. 26, pp. 16766–16769, 2002.

[37] V. Kralj-Iglič, S. Svetina, and B. ŽekŽ, “Shapes of bilayer vesicles with membrane embedded molecules,” European biophysics journal, vol. 24, no. 5, pp. 311–321, 1996.

[38] V. Kralj-Iglič, G. Pocsfalvi, L. Mesarec, V. ŠuŠtar, H. Hägerstrand, and A. Iglič, “Minimizing isotropic and deviatoric membrane energy–an unifying formation mechanism of different cellular membrane nanovesicle types,” PloS one, vol. 15, no. 12, p. e0244796, 2020.

[39] U. Seifert, “Configurations of fluid membranes and vesicles,” Advances in physics, vol. 46, no. 1, pp. 13–137, 1997.

[40] D. Kabaso, R. Shlomovitz, T. Auth, V. L. Lew, and N. S. Gov, “Curling and local shape changes of red blood cell membranes driven by cytoskeletal reorganization,” Biophysical journal, vol. 99, no. 3, pp. 808–816, 2010.

[41] R. Lipowsky, “Spontaneous tubulation of membranes and vesicles reveals membrane tension generated by spontaneous curvature,” Faraday discussions, vol. 161, pp. 305–331, 2013.

[42] R. Lipowsky and H.-G. Döbereiner, “Vesicles in contact with nanoparticles and colloids,” EPL (Europhysics Letters), vol. 43, no. 2, p. 219, 1998.

[43] M. Deserno and T. Bickel, “Wrapping of a spherical colloid by a fluid membrane,” EPL (Europhysics Letters), vol. 62, no. 5, p. 767, 2003.

[44] L. H. Miller, S. J. Mason, D. F. Clyde, and M. H. McGinniss, “The resistance factor to plasmodium vivax in blacks: the duffy-blood-group genotype, fyfy,” New England Journal of Medicine, vol. 295, no. 6, pp. 302–304, 1976.

[45] L. Foret, “Shape and energy of a membrane bud induced by protein coats or viral protein assembly,” The European Physical Journal E, vol. 37, no. 5, pp. 1–13, 2014.

[46] N. Mohandas and E. Evans, “Mechanical properties of the red cell membrane in relation to molecular structure and genetic defects,” Annual review of biophysics and biomolecular structure, vol. 23, no. 1, pp. 787–818, 1994.

[47] N. Mohandas and P. G. Gallagher, “Red cell membrane: past, present, and future,” Blood, vol. 112, no. 10, pp. 3939–3948, 2008.

[48] H. Alimohamadi, A. S. Smith, R. B. Nowak, V. M. Fowler, and P. Rangamani, “Non-uniform distribution of myosin-mediated forces governs red blood cell membrane curvature through tension modulation,” PLOS Computational Biology, vol. 16, no. 5, p. e1007890, 2020.

[49] R. B. Nowak, H. Alimohamadi, K. Pestonjamasp, P. Rangamani, and V. M. Fowler, “Nanoscale dynamics of actin filaments in the red blood cell membrane skeleton,” Molecular biology of the cell, vol. 33, no. 3, p. ar28, 2022.

[50] B. Genton, A.-Y. Fadwa, C. S. Mgone, N. Alexander, M. M. Paniu, M. P. Alpers, and D. Mokela, “Ovalocytosis and cerebral malaria,” Nature, vol. 378, no. 6557, pp. 564–565, 1995.

[51] C. Kidson, G. Lamont, A. Saul, and G. T. Nurse, “Ovalocytic erythrocytes from melanesians are resistant to invasion by malaria parasites in culture,” Proceedings of the National Academy of Sciences, vol. 78, no. 9, pp. 5829–5832, 1981.

[52] A. R. Dluzewski, G. B. Nash, R. J. Wilson, D. M. Reardon, and W. B. Gratzer, “Invasion of hereditary ovalocytes by plasmodium falciparum in vitro and its relation to intracellular atp concentration,” Molecular and biochemical parasitology, vol. 55, no. 1-2, pp. 1–7, 1992.

[53] A. Dluzewski, R. Wilson, W. Gratzer, et al., “Relation of red cell membrane properties to invasion by plasmodium falciparum,” Parasitology, vol. 91, no. 2, pp. 273–280, 1985.

[54] X. Sisquella, T. Nebl, J. K. Thompson, L. Whitehead, B. M. Malpede, N. D. Salinas, K. Rogers, N. H. Tolia, A. Fleig, J. O’Neill, et al., “Plasmodium falciparum ligand binding to erythrocytes induce alterations in deformability essential for invasion,” Elife, vol. 6, p. e21083, 2017.

[55] C. Msosa, T. Abdalrahman, and T. Franz, “An analytical model describing the mechanics of erythrocyte membrane wrapping during active invasion of a plasmodium falciparum merozoite,” Journal of the Mechanical Behavior of Biomedical Materials, vol. 140, p. 105685, 2023.

[56] D. A. Fedosov, H. Lei, B. Caswell, S. Suresh, and G. E. Karniadakis, “Multiscale modeling of red blood cell mechanics and blood flow in malaria,” PLoS Comput Biol, vol. 7, no. 12, p. e1002270, 2011.

[57] R. Skalak, A. Tozeren, R. Zarda, and S. Chien, “Strain energy function of red blood cell membranes,” Biophysical journal, vol. 13, no. 3, pp. 245–264, 1973.

[58] R. Waugh and E. A. Evans, “Thermoelasticity of red blood cell membrane,” Biophysical journal, vol. 26, no. 1, pp. 115–131, 1979.

[59] R. Mukhopadhyay, H. G. Lim, and M. Wortis, “Echinocyte shapes: bending, stretching, and shear determine spicule shape and spacing,” Biophysical Journal, vol. 82, no. 4, pp. 1756–1772, 2002.

[60] D. Kuzman, S. Svetina, R. Waugh, and B. ŽekŠ, “Elastic properties of the red blood cell membrane that determine echinocyte deformability,” European Biophysics Journal, vol. 33, no. 1, pp. 1–15, 2004.

[61] S. Svetina, D. Kuzman, R. E. Waugh, P. Ziherl, and B. ŽekŠ, “The cooperative role of membrane skeleton and bilayer in the mechanical behaviour of red blood cells,” Bioelectrochemistry, vol. 62, no. 2, pp. 107–113, 2004.

[62] S. Svetina, G. Kokot, T. Š. Kebe, B. ŽekŠ, and R. E. Waugh, “A novel strain energy relationship for red blood cell membrane skeleton based on spectrin stiffness and its application to micropipette deformation,” Biomechanics and modeling in mechanobiology, vol. 15, no. 3, pp. 745–758, 2016.

[63] Z. Feng, R. E. Waugh, and Z. Peng, “Constitutive model of erythrocyte membranes with distributions of spectrin orientations and lengths,” Biophysical Journal, vol. 119, no. 11, pp. 2190–2204, 2020.

[64] J. T. Jenkins, “The equations of mechanical equilibrium of a model membrane,” SIAM Journal on Applied Mathematics, vol. 32, no. 4, pp. 755–764, 1977.

[65] J. Jenkins, “Static equilibrium configurations of a model red blood cell,” Journal of mathematical biology, vol. 4, no. 2, pp. 149–169, 1977.

[66] B. K. Pai and H. D. Weymann, “Equilibrium shapes of red blood cells in osmotic swelling,” Journal of biomechanics, vol. 13, no. 2, pp. 105–112, 1980.

[67] R. Lipowsky, “Budding of membranes induced by intramembrane domains,” Journal de Physique II, vol. 2, no. 10, pp. 1825–1840, 1992.

[68] F. Frey and U. S. Schwarz, “Competing pathways for the invagination of clathrin-coated membranes,” Soft Matter, 2020.

[69] V. Kralj-Iglič, V. Heinrich, S. Svetina, and B. ŽekŠ, “Free energy of closed membrane with anisotropic inclusions,” The European Physical Journal B-Condensed Matter and Complex Systems, vol. 10, no. 1, pp. 5–8, 1999.

[70] P. B. Canham, “The minimum energy of bending as a possible explanation of the biconcave shape of the human red blood cell,” Journal of theoretical biology, vol. 26, no. 1, pp. 61–81, 1970.

[71] S. K. Boey, D. H. Boal, and D. E. Discher, “Simulations of the erythrocyte cytoskeleton at large deformation. microscopic models,” Biophysical Journal, vol. 75, no. 3, pp. 1573–1583, 1998.

[72] D. E. Discher, D. H. Boal, and S. K. Boey, “Simulations of the erythrocyte cytoskeleton at large deformation. micropipette aspiration,” Biophysical Journal, vol. 75, no. 3, pp. 1584–1597, 1998.

[73] J. F. Marko and E. D. Siggia, “Stretching dna,” Macromolecules, vol. 28, no. 26, pp. 8759–8770, 1995.

[74] C. Bustamante, Z. Bryant, and S. B. Smith, “Ten years of tension: single-molecule dna mechanics,” Nature, vol. 421, no. 6921, pp. 423–427, 2003.

[75] B. Hendrickson, M. Shirani, and D. J. Steigmann, “Equilibrium theory for a lipid bilayer with a conforming cytoskeletal membrane,” Mathematics and Mechanics of Complex Systems, vol. 8, no. 1, pp. 69–99, 2020.

[76] A. Iglič, “A possible mechanism determining the stability of spiculated red blood cells,” Journal of biomechanics, vol. 30, no. 1, pp. 35–40, 1997.

[77] A. Iglič, V. Kralj-Iglič, and H. Hägerstrand, “Stability of spiculated red blood cells induced by intercalation of amphiphiles in cell membrane,” Medical and Biological Engineering and Computing, vol. 36, no. 2, pp. 251–255, 1998.

[78] R. E. Waugh, “Elastic energy of curvature-driven bump formation on red blood cell membrane,” Biophysical journal, vol. 70, no. 2, pp. 1027–1035, 1996.

[79] E. Evans, “New membrane concept applied to the analysis of fluid shear-and micropipette-deformed red blood cells,” Biophysical journal, vol. 13, no. 9, pp. 941–954, 1973.

[80] U. Seifert and R. Lipowsky, “Adhesion of vesicles,” Physical Review A, vol. 42, no. 8, p. 4768, 1990.

[81] A. Agrawal and D. J. Steigmann, “Coexistent fluid-phase equilibria in biomembranes with bending elasticity,” Journal of Elasticity, vol. 93, no. 1, pp. 63–80, 2008.

[82] H. Alimohamadi, M. K. Bell, S. Halpain, and P. Rangamani, “Mechanical principles governing the shapes of dendritic spines,” Frontiers in Physiology, vol. 12, 2021.

[83] N. Walani, J. Torres, and A. Agrawal, “Endocytic proteins drive vesicle growth via instability in high membrane tension environment,” Proceedings of the National Academy of Sciences, vol. 112, no. 12, pp. E1423–E1432, 2015.

[84] J. Li, M. Dao, C. Lim, and S. Suresh, “Spectrin-level modeling of the cytoskeleton and optical tweezers stretching of the erythrocyte,” Biophysical journal, vol. 88, no. 5, pp. 3707–3719, 2005.

[85] L. Pan, R. Yan, W. Li, and K. Xu, “Super-resolution microscopy reveals the native ultrastructure of the erythrocyte cytoskeleton,” Cell reports, vol. 22, no. 5, pp. 1151–1158, 2018.

[86] H. Alimohamadi, R. Vasan, J. Hassinger, J. C. Stachowiak, and P. Rangamani, “The role of traction in membrane curvature generation,” Mol. Biol. Cell, vol. 29, no. 16, pp. 2024–2035, 2018.

[87] H. Alimohamadi, B. Ovryn, and P. Rangamani, “Modeling membrane nanotube morphology: the role of heterogeneity in composition and material properties,” Scientific Reports, vol. 10, no. 1, pp. 1–15, 2020.

[88] R. Vasan, S. Rudraraju, M. Akamatsu, K. Garikipati, and P. Rangamani, “A mechanical model reveals that non-axisymmetric buckling lowers the energy barrier associated with membrane neck constriction,” Soft Matter, vol. 16, no. 3, pp. 784–797, 2020.

[89] C. Zhu, C. T. Lee, and P. Rangamani, “Mem3dg: modeling membrane mechanochemical dynamics in 3d using discrete differential geometry,” Biophysical reports, vol. 2, no. 3, p. 100062, 2022.

[90] D. Auddya, X. Zhang, R. Gulati, R. Vasan, K. Garikipati, P. Rangamani, and S. Rudraraju, “Biomembranes undergo complex, non-axisymmetric deformations governed by kirchhoff–love kinematicsand revealed by a three-dimensional computational framework,” Proceedings of the Royal Society A, vol. 477, no. 2255, p. 20210246, 2021.

[91] H. Alimohamadi, Application of continuum mechanics for a variety of curvature generation phenomena in cell biophysics. University of California, San Diego, 2021.

[92] M. Chabanon and P. Rangamani, “Gaussian curvature directs the distribution of spontaneous curvature on bilayer membrane necks,” Soft Matter, 2018.

[93] M. Chabanon and P. Rangamani, “Geometric coupling of helicoidal ramps and curvature-inducing proteins in organelle membranes,” Journal of the Royal Society Interface, vol. 16, no. 158, p. 20190354, 2019.

[94] J. Evans, W. Gratzer, N. Mohandas, K. Parker, and J. Sleep, “Fluctuations of the red blood cell membrane: relation to mechanical properties and lack of atp dependence,” Biophysical journal, vol. 94, no. 10, pp. 4134–4144, 2008.

[95] G. Popescu, T. Ikeda, K. Goda, C. A. Best-Popescu, M. Laposata, S. Manley, R. R. Dasari, K. Badizadegan, and M. S. Feld, “Optical measurement of cell membrane tension,” Physical review letters, vol. 97, no. 21, p. 218101, 2006.

[96] A. J. Crick, M. Theron, T. Tiffert, V. L. Lew, P. Cicuta, and J. C. Rayner, “Quantitation of malaria parasite-erythrocyte cell-cell interactions using optical tweezers,” Biophysical journal, vol. 107, no. 4, pp. 846–853, 2014.

[97] J. Agudo-Canalejo and R. Lipowsky, “Critical particle sizes for the engulfment of nanoparticles by membranes and vesicles with bilayer asymmetry,” ACS nano, vol. 9, no. 4, pp. 3704–3720, 2015.

[98] H. Schönherr, J. M. Johnson, P. Lenz, C. W. Frank, and S. G. Boxer, “Vesicle adsorption and lipid bilayer formation on glass studied by atomic force microscopy,” Langmuir, vol. 20, no. 26, pp. 11600–11606, 2004.

[99] T. Gruhn, T. Franke, R. Dimova, and R. Lipowsky, “Novel method for measuring the adhesion energy of vesicles,” Langmuir, vol. 23, no. 10, pp. 5423–5429, 2007.

[100] T. H. Anderson, Y. Min, K. L. Weirich, H. Zeng, D. Fygenson, and J. N. Israelachvili, “Formation of supported bilayers on silica substrates,” Langmuir, vol. 25, no. 12, pp. 6997–7005, 2009.

[101] J. Liu, M. Kaksonen, D. G. Drubin, and G. Oster, “Endocytic vesicle scission by lipid phase boundary forces,” Proceedings of the National Academy of Sciences, vol. 103, no. 27, pp. 10277–10282, 2006.

[102] A. J. García-Sáez, S. Chiantia, and P. Schwille, “Effect of line tension on the lateral organization of lipid membranes,” Journal of Biological Chemistry, vol. 282, no. 46, pp. 33537–33544, 2007.

[103] A. Swihart, J. Mikrut, J. Ketterson, and R. Macdonald, “Atomic force microscopy of the erythrocyte membrane skeleton,” Journal of microscopy, vol. 204, no. 3, pp. 212–225, 2001.

[104] M. Takeuchi, H. Miyamoto, Y. Sako, H. Komizu, and A. Kusumi, “Structure of the erythrocyte membrane skeleton as observed by atomic force microscopy,” Biophysical Journal, vol. 74, no. 5, pp. 2171–2183, 1998.

[105] J. A. Ursitti, D. W. Pumplin, J. B. Wade, and R. J. Bloch, “Ultrastructure of the human erythrocyte cytoskeleton and its attachment to the membrane,” Cell motility and the cytoskeleton, vol. 19, no. 4, pp. 227–243, 1991.

[106] J. A. Ursitti and J. B. Wade, “Ultrastructure and immunocytochemistry of the isolated human erythrocyte membrane skeleton,” Cell motility and the cytoskeleton, vol. 25, no. 1, pp. 30–42, 1993.

[107] T. J. Byers and D. Branton, “Visualization of the protein associations in the erythrocyte membrane skeleton,” Proceedings of the National Academy of Sciences, vol. 82, no. 18, pp. 6153–6157, 1985.

[108] S.-C. Liu, L. H. Derick, and J. Palek, “Visualization of the hexagonal lattice in the erythrocyte membrane skeleton.,” The Journal of cell biology, vol. 104, no. 3, pp. 527–536, 1987.

[109] A. M. McGough and R. Josephs, “On the structure of erythrocyte spectrin in partially expanded membrane skeletons,” Proceedings of the National Academy of Sciences, vol. 87, no. 13, pp. 5208–5212, 1990.

[110] A. H. Bahrami, “Orientational changes and impaired internalization of ellipsoidal nanoparticles by vesicle membranes,” Soft Matter, vol. 9, no. 36, pp. 8642–8646, 2013.

[111] F. Frey, F. Ziebert, and U. S. Schwarz, “Dynamics of particle uptake at cell membranes,” Physical Review E, vol. 100, no. 5, p. 052403, 2019.

[112] E. A. Evans, Mechanics and thermodynamics of biomembranes. CRC press, 2018.

[113] S. Henon, G. Lenormand, A. Richert, and F. Gallet, “A new determination of the shear modulus of the human erythrocyte membrane using optical tweezers,” Biophysical journal, vol. 76, no. 2, pp. 1145–1151, 1999.

[114] G. Lenormand, S. Hénon, A. Richert, J. Siméon, and F. Gallet, “Direct measurement of the area expansion and shear moduli of the human red blood cell membrane skeleton,” Biophysical journal, vol. 81, no. 1, pp. 43–56, 2001.

[115] N. D. Geoghegan, C. Evelyn, L. W. Whitehead, M. Pasternak, P. McDonald, T. Triglia, D. S. Marapana, D. Kempe, J. K. Thompson, M. J. Mlodzianoski, et al., “4d analysis of malaria parasite invasion offers insights into erythrocyte membrane remodeling and parasitophorous vacuole formation,” Nature communications, vol. 12, no. 1, p. 3620, 2021.

[116] J. Hansen, R. Skalak, S. Chien, and A. Hoger, “An elastic network model based on the structure of the red blood cell membrane skeleton,” Biophysical journal, vol. 70, no. 1, pp. 146–166, 1996.

[117] T. G. Fai, A. Leo-Macias, D. L. Stokes, and C. S. Peskin, “Image-based model of the spectrin cytoskeleton for red blood cell simulation,” PLoS computational biology, vol. 13, no. 10, p. e1005790, 2017.

[118] M. Dao, J. Li, and S. Suresh, “Molecularly based analysis of deformation of spectrin network and human erythrocyte,” Materials Science and Engineering: C, vol. 26, no. 8, pp. 1232–1244, 2006.

[119] N. Gov and S. Safran, “Red blood cell membrane fluctuations and shape controlled by atp-induced cytoskeletal defects,” Biophysical journal, vol. 88, no. 3, pp. 1859–1874, 2005.

[120] K. Ayi, W. C. Liles, P. Gros, and K. C. Kain, “Adenosine triphosphate depletion of erythrocytes simulates the phenotype associated with pyruvate kinase deficiency and confers protection against plasmodium falciparum in vitro,” The Journal of infectious diseases, vol. 200, no. 8, pp. 1289–1299, 2009.

[121] Y. Park, C. A. Best, T. Auth, N. S. Gov, S. A. Safran, G. Popescu, S. Suresh, and M. S. Feld, “Metabolic remodeling of the human red blood cell membrane,” Proceedings of the National Academy of Sciences, vol. 107, no. 4, pp. 1289–1294, 2010.

[122] A. Ghisleni, M. Bonilla-Quintana, M. Crestani, A. Fukuzawa, P. Rangamani, and N. Gauthier, “Mechanically induced topological transition of spectrin regulates its distribution in the mammalian cortex,” bioRxiv, pp. 2023–01, 2023.

[123] H. Turlier, D. A. Fedosov, B. Audoly, T. Auth, N. S. Gov, C. Sykes, J.-F. Joanny, G. Gompper, and T. Betz, “Equilibrium physics breakdown reveals the active nature of red blood cell flickering,” Nature physics, vol. 12, no. 5, pp. 513–519, 2016.

[124] A. F. Cowman and B. S. Crabb, “Invasion of red blood cells by malaria parasites,” Cell, vol. 124, no. 4, pp. 755–766, 2006.

[125] S. Hegge, K. Uhrig, M. Streichfuss, G. Kynast-Wolf, K. Matuschewski, J. P. Spatz, and F. Frischknecht, “Direct manipulation of malaria parasites with optical tweezers reveals distinct functions of plasmodium surface proteins,” ACS nano, vol. 6, no. 6, pp. 4648–4662, 2012.

[126] A. F. Cowman, J. Healer, D. Marapana, and K. Marsh, “Malaria: biology and disease,” Cell, vol. 167, no. 3, pp. 610–624, 2016.

[127] J. Langhorne, F. M. Ndungu, A.-M. Sponaas, and K. Marsh, “Immunity to malaria: more questions than answers,” Nature immunology, vol. 9, no. 7, pp. 725–732, 2008.

[128] P. C. Sabeti, D. E. Reich, J. M. Higgins, H. Z. Levine, D. J. Richter, S. F. Schaffner, S. B. Gabriel, J. V. Platko, N. J. Patterson, G. J. McDonald, et al., “Detecting recent positive selection in the human genome from haplotype structure,” Nature, vol. 419, no. 6909, pp. 832–837, 2002.

[129] A. Ciana, C. Achilli, and G. Minetti, “Membrane rafts of the human red blood cell,” Molecular membrane biology, vol. 31, no. 2-3, pp. 47–57, 2014.

[130] S. C. Murphy, B. U. Samuel, T. Harrison, K. D. Speicher, D. W. Speicher, M. E. Reid, R. Prohaska, P. S. Low, M. J. Tanner, N. Mohandas, et al., “Erythrocyte detergent-resistant membrane proteins: their characterization and selective uptake during malarial infection,” Blood, vol. 103, no. 5, pp. 1920–1928, 2004.

[131] A. George, S. Pushkaran, L. Li, X. An, Y. Zheng, N. Mohandas, C. H. Joiner, and T. A. Kalfa, “Altered phosphorylation of cytoskeleton proteins in sickle red blood cells: the role of protein kinase c, rac gtpases, and reactive oxygen species,” Blood Cells, Molecules, and Diseases, vol. 45, no. 1, pp. 41–45, 2010.

